# Sensing cellular growth rate facilitates its robust optimal adaptation to changing conditions

**DOI:** 10.1101/2024.07.09.602663

**Authors:** Robert Planqué, Josephus Hulshof, Frank J. Bruggeman

## Abstract

The determinants of growth rate and the associated metabolism has been at center stage in microbial physiology for over seventy years. In this paper we show that a cell sensing its own growth rate is in principle capable of maximising it using a gene regulatory circuit responsible for adapting metabolic enzyme concentrations in dynamic conditions. This is remarkable, since any state of (close-to) optimal growth depends on nutrient conditions, and is thus not a fixed target. We derive the properties of such gene regulatory networks, and prove that such circuits allow the growth rate to be a Lyapunov function. We derive this from a general stoichiometric and kinetic description of cellular metabolism. Interestingly, our finding is in agreement with our current understanding of how *E. coli* controls its growth rate. It uses ppGpp to tune the growth rate by balancing metabolic and ribosomal protein expression. Since ppGpp covaries 1-to-1 with the protein translation rate, an excellent proxy for growth rate, on a timescale of seconds, this suggests that direct sensing of the growth rate underlies growth rate optimisation in *E. coli*.

## Introduction

Microorganisms exhibit an extraordinary spectrum of biodiversity, evident in the myriad environmental conditions they inhabit [19]. Despite this diversity, the underlying metabolism – its kinetic, regulatory and bioenergetic principles – can likely be understood in terms of a common set of principles [39, 7, 55, 5].

One fundamental concept contributing to this understanding is the “unity of biochemistry.” This encompasses the evolutionary conservation of enzyme kinetics, the allosteric regulation of enzyme catalysis, cellular bioenergetics, and the basic organization of cellular metabolism into energy- and precursor-producing catabolic pathways driving the biosynthesis of macromolecules [39].

Additionally, the principle of natural selection plays a pivotal role [16, 9]. Fit genotypes, those that produce more offspring, prevail over those that over time reproduced less. Achieving a competitive time-averaged growth rate requires microbial cells to allocate resources judiciously, balancing protein-demanding tasks crucial for adaptation, stress tolerance, and growth to enhance their fitness [6, 55].

These principles hold true across microbial species and environmental niches, underscoring their universal significance in microbial physiology. Advancements in systems biology over the past three decades have propelled this understanding forward, particularly through the development of genome-scale models of metabolic networks [47] and the development of the concepts of finite biosynthetic resources [36, 55]. However, these contributions have primarily focused on balanced growth conditions [50]; theoretical principles, modeling methodologies, and the integration of “omics” data have mostly been developed for exponentially growing cultures displaying steady state metabolism [17, 42].

We still lack an understanding in fundamental terms of the dynamic process of adaptation to new conditions [35, 18]. Do general principles of gene-expression regulation by environmental sensing exist? In particular, upon an environmental change, the growth rate generally drops; sensing and gene expression controls are activated, cells adjust their metabolic activity by adjusting protein concentrations, and improve their growth rate [7]. The theoretical principles for balanced growth conditions allow us to deduce from first principles (genomic information, enzyme kinetics, flux balance, finite biosynthetic resource allocation, evolutionary theory) which metabolic strategy achieves optimal balanced growth in which conditions—it provides predictions of the end points of the adaptation process [36, 61, 38, 56, 60, 12, 11]. Our aim here is to build a similar framework for dynamical gene expression regulation for growth-rate adaptation, again from first principles. Pioneering studies do exist, and inspired this work [3, 18, 33, 45, 62, 8, 15].

We address how a microbial cell that needs to adapt to a new condition should control its gene expression in order to attain a competitive growth rate. This competitive growth rate is bounded by the growth rate that is maximally achievable given the organism’s genetic potential and the prevailing environmental conditions. We consider realistic biochemical models of cellular adaptation that incorporate metabolism, growth and gene expression, involving a gene regulatory network that uses a read-out of the physiological state as input, and induces protein synthesis rates as output, which are eventually balanced by ‘dilution by growth’ when a steady-state of balanced growth has settled [45]. Coupling the metabolic network and gene expression network establishes a single dynamic system in which environmental changes modify the physiological state through metabolism; a readout of this state induces changes in gene expression, leading to new protein concentrations, which are fed back into metabolism, resulting in a self-organising dynamics.

In this paper, we show that for the problem of optimal adaptation of growth rate to changing conditions, the action of the gene network can be derived from first principles. It turns out that the biochemical description of the metabolic network uniquely determines it. Importantly, when the instantaneous growth rate is chosen as physiological input variable to this gene network, we prove that the growth rate itself becomes a Lyapunov function and therefore always increases in time, consistent with much experimental data on bacterial growth in nutrient up- and downshifts [18, 37, 33, 65, 40, 44].

Lyapunov functions play an important role in dynamical systems theory and control theory to show that steady states of systems are (asymptotically, or even globally) stable or reachable by control mechanisms [25, 20, 26]. Here we exploit this same idea: by showing that the growth rate is a Lyapunov function, we show that the coupled metabolic-enzyme system always reaches a steady state of maximal growth, regardless of the environmental state. This implies that sensing any system variable that is in one-to-one correspondence with the instantaneous growth rate would enable the cell to adjust its gene expression and navigate in the direction of an increasing growth rate, thus enhancing its competitive potential.

Recent experimental work [62] suggests that *Escherichia coli* appears to control its growth rate according to this principle. The concentration of the regulatory molecule ppGpp appears to be a linear function of the reciprocal of the ribosomal translation rate in both steady state and dynamic conditions [62, 64], and to covary with steady state cellular growth rate [62, 49]. Thus, the growth rate is sensed by ppGpp. Since ppGpp is responsible for regulating the balance between adaptation, stress and growth-associated protein expression via its effects on the expressioon of ribosomal, catabolic and sigma-factor genes [46], *E. coli* appears to sense its growth rate to adjust itself to new conditions. Here we rationalise this by showing that exactly for the task of judicious allocation of biosynthetic resources in changing conditions the growth rate is a Lyapunov function, making it the perfect guide for navigating dynamic environments.

## Results

### Introducing the framework of a self-replicating microbial cell

In Figure 1 we introduce a simplified model of metabolism and gene expression that contains all the ingredients to understand our main result. This type of coarse-grained model is widely used in the literature [36, 60, 13, 18, 56, 4, 8]. The mathematical description of more detailed models is given in the Methods.

**Figure 1.**
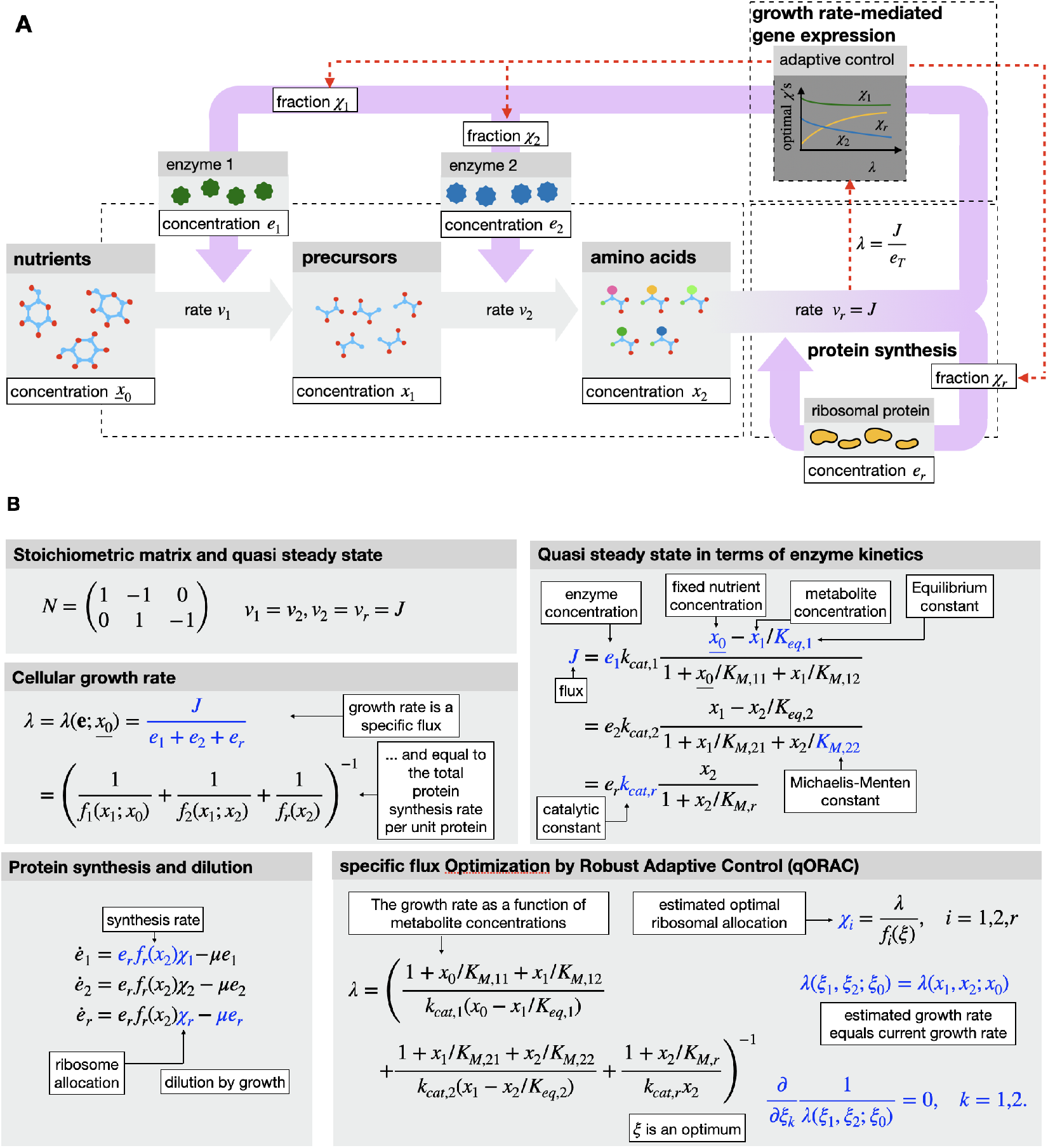
The essence of the coupling of the metabolism to the synthesis of proteins and the regulation of protein expression in a self-replicating microbial cell. **A**. A schematic overview of metabolism. The synthesis of amino acids from nutrients by enzymes is shown, whose concentrations are determined by the fraction of ribosomes allocated to their synthesis (in proportion to the transcript (mRNA) fraction of the enzymes). The genetic control network (not shown) regulates the ribosomal fraction such that the enzymes attain their (close-to-)optimal values. We assume that the metabolic network operates at quasi-steady state relative to the genetic network, which is dynamic. **B**. The essential aspects of the modeling formalism.

We consider a nutrient with concentration *<u>x</u>*_<u>0</u>_ in the environment and assumed constant. It is taken up by the cell and converted into precursor molecules by a first enzyme at a rate *v*_1_. We assume that this rate is given by *v*_1_ = *e*_1_*f*_1_(*x*_1_; *<u>x</u>*_<u>0</u>_), where *x*_1_ is the concentration of precursors, *e*_1_ the concentration of enzyme 1, and *f*_1_ a nonlinear, kinetic rate law [10], detailing the dependence of the reaction rate on concentration of substrates, products and kinetic parameters (including the reaction’s equilibrium constant and catalytic rate constant *k*_*cat*,1_).

The precursors are subsequently converted into amino acids (concentration *x*_2_) by a second enzyme with concentration *e*_2_ and at a rate *v*_2_ = *e*_2_*f*_2_(*x*_1_, *x*_2_). The amino acids are finally translated into protein by ribosomes at a rate *v*_*r*_ = *e*_*r*_*f*_*r*_(*x*_2_). Since the ribosome synthesizes all the different kinds of proteins, it needs to be allocated accordingly. Let *χ*_1_ be the fraction of actively translating ribosomes allocated to synthesis of enzyme 1, then enzyme 1 is produced at a rate *χ*_1_*v*_*r*_, and similarly for enzyme 2 and the ribosome itself; then *χ*_1_ + *χ*_2_ + *χ*_*r*_ = 1.

As a matter of notation, we use ***x*** to denote the internal metabolite concentrations (*x*_1_, *x*_2_), viewing *<u>x</u>*_<u>0</u>_ as a parameter. We sometimes abbreviate the kinetic rate laws by *f*_*i*_(***x***; *<u>x</u>*_<u>0</u>_). The context should make clear on which variables and parameters *f*_*i*_ actually depends. Figure 1B provides an explicit choice of the kinetic rate laws used in the coarse grained model.

#### Enzyme dynamics, quasi steady state metabolism and cellular growth rate

Enzyme and ribosome concentrations change because of the imbalance between their rates of biosynthesis and dilution by growth,

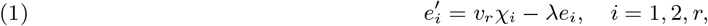

where *λ* is the cellular growth rate. Assuming that the total protein concentration is constant,

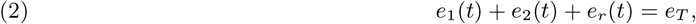

equation (1) indicates a definition of the (steady-state or balanced) growth rate as the protein translation rate per unit cellular protein [13, 11, 14, 54],

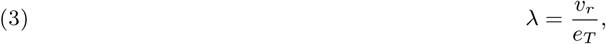

which makes the growth rate a specific flux (flux per unit enzyme concentration). Substituting eq. (3) into eq. (1) gives

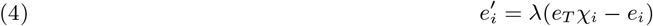

showing that at steady state the ribosomal fraction synthesizing enzyme *i* is equal to the corresponding protein fraction

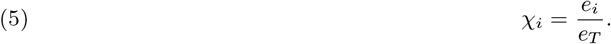

We will thus interchange freely between “ribosome allocation” and “enzyme allocation” as they are essentially one and the same.

We further assume that changes in protein concentrations occur at time scales that are significantly longer than metabolic changes. In other words, we assume that metabolism is in so-called quasi steady state with respect to changes in protein concentrations. For specific values of *<u>x</u>*_<u>0</u>_, *e*_1_, *e*_2_ and *e*_*r*_, the internal quasi steady state metabolite concentrations are fixed.

In this state, metabolic mass flows are balanced, so *v*_1_ = *v*_2_ = *v*_*r*_. Since *v*_*r*_ and *λ* are so tightly coupled by eq. (3), we choose *v*_*r*_ to be *the* (quasi steady state) flux through the metabolic pathway, and denote it by *J*, so that eq. (3) now reads

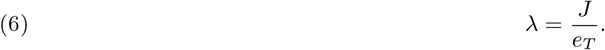

Any quasi steady state flux vector (*v*_1_, *v*_2_, *v*_*r*_)^*T*^ is thus of the form *J* (1, 1, 1)^*T*^. This makes our simplified model an elementary metabolic network [52, 53]. These are “one flux degree of freedom” networks [58] (choosing one flux value determines all flux values) that arise naturally as the subnetworks within larger networks that maximise growth rate in constant conditions [61, 38, 11, see Methods].

Since *v*_*i*_ = *e*_*i*_*f*_*i*_(***x***) = *J* in quasi steady state, *e*_*i*_*/J* = 1*/f*_*i*_(***x***), and [41, 45, 14]

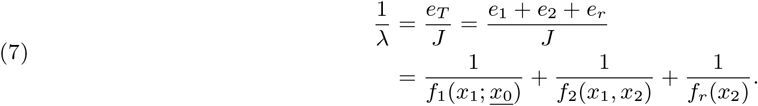

There exist therefore various ways of viewing the steady-state growth rate in a given nutrient condition:

- as the total protein synthesis rate per unit protein, cf. eq. (3) and (6);
- as the outcome of ribosome allocation, *λ* = *λ*(***χ***; *<u>x</u>*_<u>0</u>_);
- as the outcome of protein expression, *λ* = *λ*(***e***; *<u>x</u>*_<u>0</u>_);
- as a function of internal quasi steady state metabolite concentrations, cf. eq. (7), so *λ* = *λ*(***x***; *<u>x</u>*_<u>0</u>_).

(We abuse notation somewhat here, and denote all growth rate functions by the same *λ*.) We will need all these different viewpoints below.

### The essentials of a robust growth-rate maximising gene network

We now proceed to discuss competitive growth rates. We consider the extreme case of a genetic network, regulating the expression of catabolic and anabolic enzymes, that is capable of maximising the growth rate. Since any real biological network always performs below or equal to this performance, this truly optimal network shall be our benchmark case. Figures 2, 3 and 4 aid in the explanation below.

**Figure 2.**
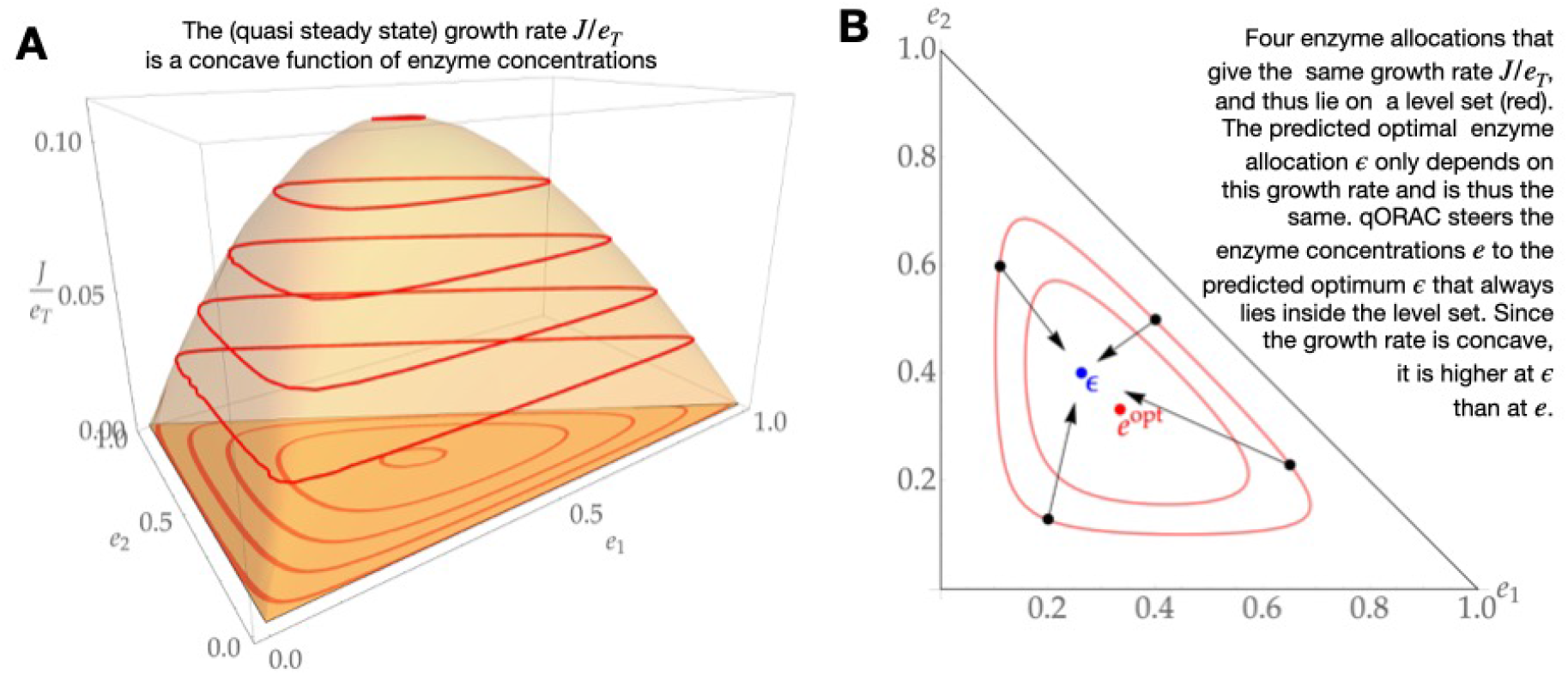
Concavity and control of specific flux. A: Growth rate *λ* for a short linear pathway with three reactions (see Fig. 1A) is a concave function of enzyme investment. Since each reaction is essential to sustain a flux through the pathway, the growth rate is zero if one of the enzyme concentrations is zero. Red level sets indicate enzyme allocations with the same growth rate. When projected onto the enzyme plane (darker orange) {*e*_1_, *e*_2_ ≥ 1, *e*_1_ + *e*_2_ ≤ 1}, the level sets are all convex and nested around the maximum *e*^*opt*^ (shown as a red dot in B). B: see next to figure.

**Figure 3.**
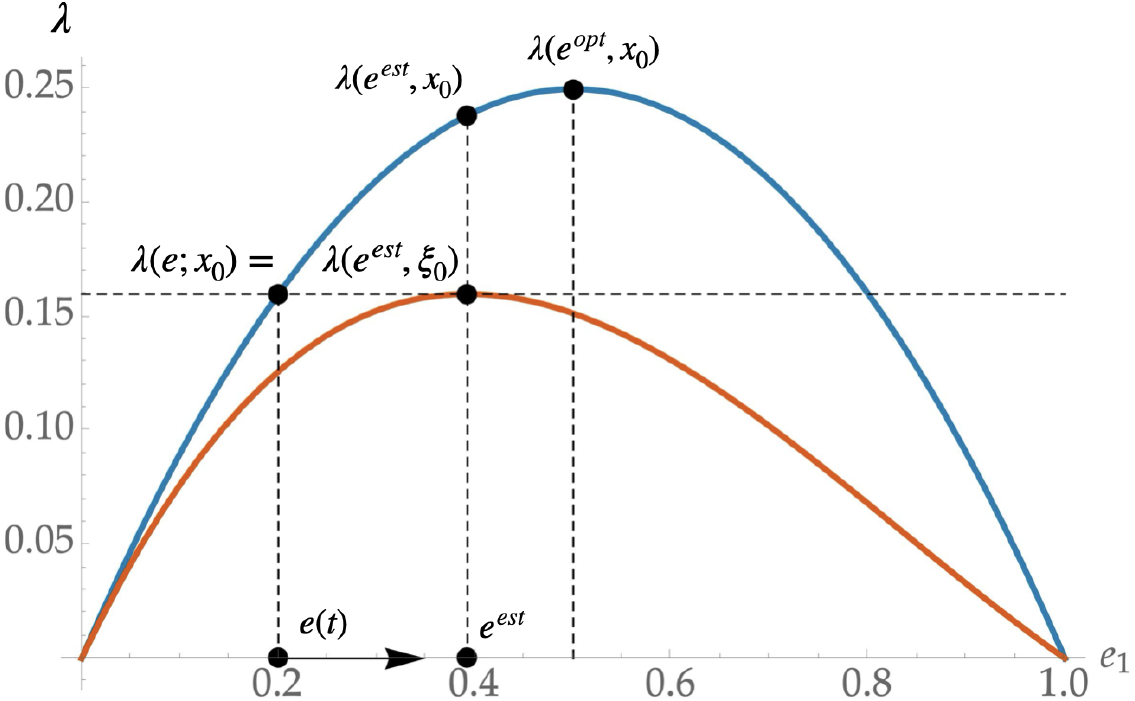
The construction of the adaptive control. We consider a pathway with just two enzymes, with *e*_1_ + *e*_2_ = 1. Consider the current set of enzyme concentrations ***e***(*t*) on the *e*_1_-axis. The growth rate at nutrient conditions *<u>x</u>*_<u>0</u>_ is indicated right above it. The adaptive control gives the optimal enzyme concentrations ***e***^*est*^ corresponding to ***e***, by choosing the unique *ξ*_0_ *< <u>x</u>*_<u>0</u>_ such that the optimal growth is equal to the current growth rate (red curve). Since *λ*(***e***; *<u>x</u>*_<u>0</u>_) is a concave function of ***e***, the position of ***e***^*est*^ is invariably such that when this estimated optimal allocation is used in the pathway with the original nutrient condition *<u>x</u>*_<u>0</u>_, a higher growth rate is achieved: *λ*(***e***^*est*^; *<u>x</u>*_<u>0</u>_) *> λ*(***e***; *<u>x</u>*_<u>0</u>_). The enzyme concentrations evolve towards the estimator (indicated by the arrow), so that the growth rate always increases. Importantly, the prediction step does not involve the nutrient concentration, but only the current growth rate. In this way, the controlled pathway robustly achieves maximal growth rate in any nutrient condition.

**Figure 4.**
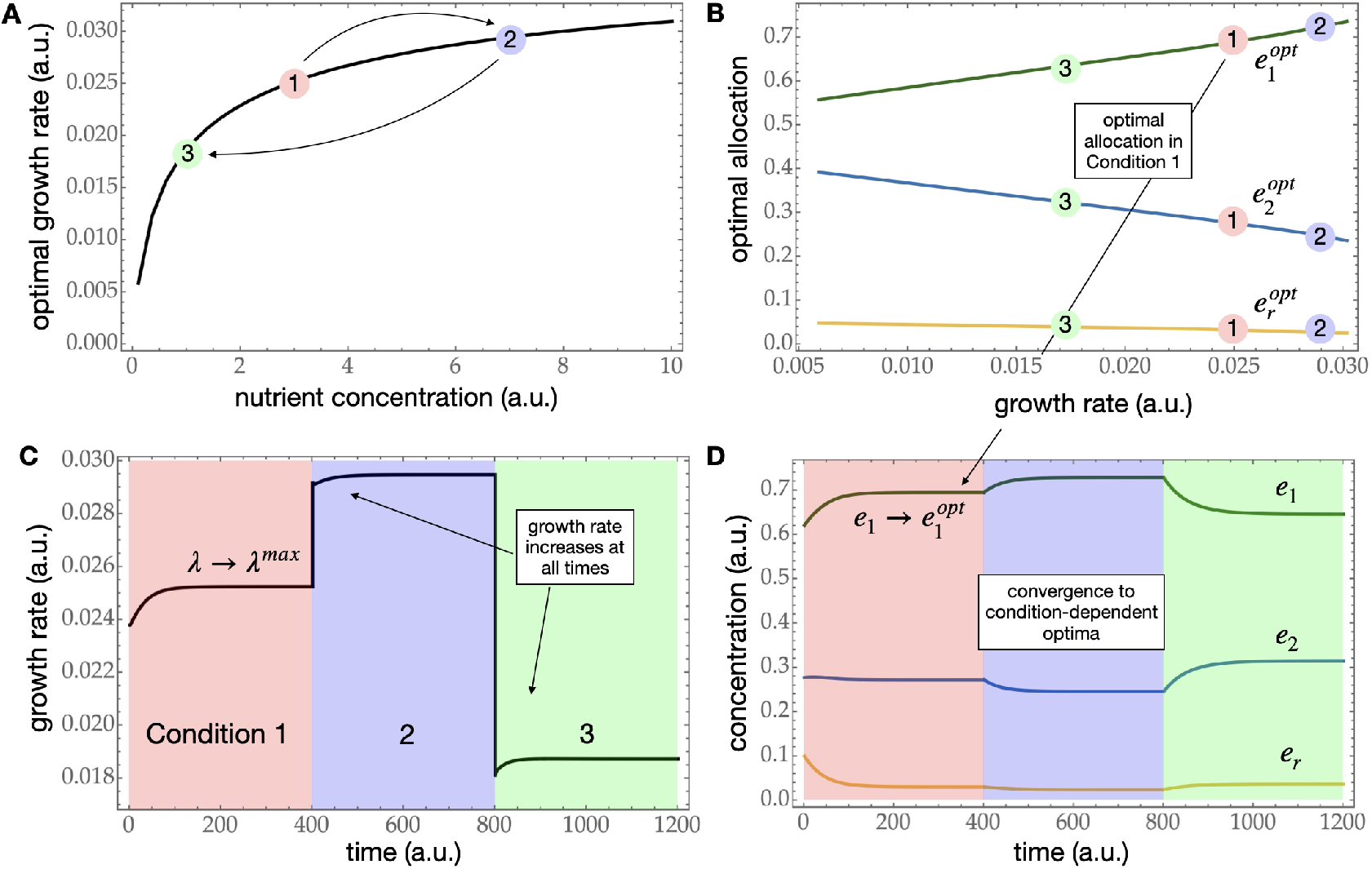
Example simulations for the adaptively controlled pathway shown in Figure 1. A: optimal growth rate, as a function of nutrient concentration. The three vertical lines correspond to the three nutrient conditions *<u>x</u>*_<u>0</u>_ = 1, 3, 7. B: the adaptive control: optimal allocation of the ribosome over the three protein synthesis reactions, as a function of growth rate. C: growth rate over time, in the three conditions shown in C. Note that the growth rate optimizes each time (compare with C). D: enzyme and ribosome concentrations over time, showing convergence to new steady state levels in each nutrient condition. The full model equations are specified in Box 1. Parameters are given in the SI.

#### In each environment there is a unique set of optimal enzyme concentrations and an optimal ribosome allocation

For any environmental condition *<u>x</u>*_<u>0</u>_, there exists a unique set of enzyme concentrations at which the steady state growth rate through the simplified network in Figure 1 is maximal. The reason is that for any reasonable choice of kinetic rate laws, such as the ones detailed in Figure 1B, relation (7) is convex in logarithmic metabolite concentrations log *x*_1_ and log *x*_2_ [41, 45]. For any *<u>x</u>*_<u>0</u>_, the optimal steady state enzyme concentrations may be found using eq. (7), by first solving

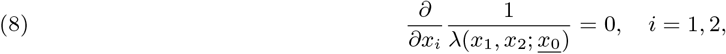

on the domain of metabolite concentrations where *f*_*i*_(***x***; *<u>x</u>*_<u>0</u>_) *>* 0, to obtain the optimal metabolite concentrations 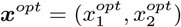, and subsequently setting (using *e*_*i*_*f*_*i*_(***x***) = *J* = *e*_*T*_ *λ*)

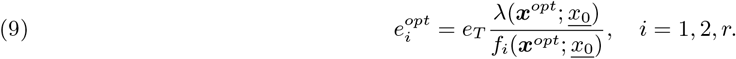

In other words,

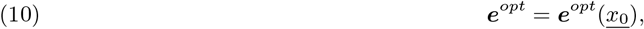

and the maximal steady state growth rate is thus in the end determined by *<u>x</u>*_<u>0</u>_ alone,

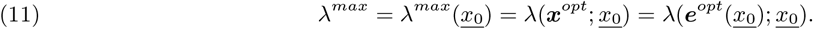

From relation (5), we conclude that the optimal steady state ribosome allocation is similarly defined by

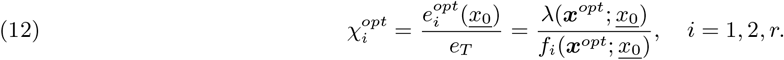

#### Increasing the nutrient concentration increases the (maximal) growth rate

For the example in Figure 1A and at fixed enzyme concentrations, the growth rate increases with increasing nutrient concentration,

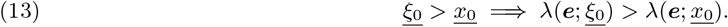

The maximal growth rate that can be attained in a certain nutrient environment therefore also increases with nutrient concentration,

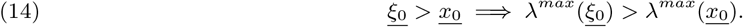

We prove this in the Supplemental Information (SI).

#### Estimating the optimal ribosome allocation using the growth rate

Equations (10), (11), and (14) may be combined to conclude that all possible optimal enzyme concentration vectors may be parameterised by the corresponding maximal growth rate,

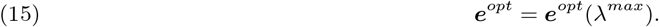

The current, instantaneous (suboptimal) growth rate *λ*(*t*) may thus be used to *estimate* the optimal enzyme concentrations by replacing *λ*^*max*^ with *λ* in eq. (15). We denote this estimate by

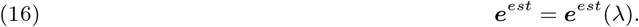

As ribosomal allocations and protein fractions are the same in steady state, cf. eq. (5), we may also introduce

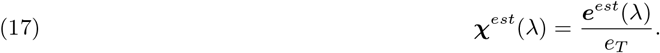

For the linear pathway in Figure 1, the dependence of this allocation on the growth rate is shown schematically in the small grey box on top of the protein translation reaction in Figure 1A.

Importantly, at steady state the estimated optimal allocation only coincides with the actual optimal allocation if and only if the growth rate is maximal (for the current nutrient condition), i.e.,

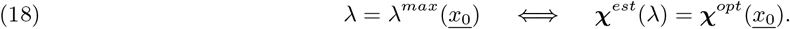

#### In suboptimal states, the growth rate may be improved by redistributing enzyme concentrations

So far we have only explored the properties of the static problem of finding optimal enzyme concentrations for a given nutrient condition. We concluded that these optimal concentrations are generally uniquely determined, and that the maximal growth rate is as well. Higher nutrient concentrations leads to a higher maximal growth rate, so the optimal allocation may be estimated from the growth rate alone, rather than from the nutrient concentration.

We now slowly turn to the problem of getting to this optimal state. First we consider the overall behaviour of the growth rate, as a function of enzyme investment, in one nutrient condition (Figure 2). As a first observation, we know that not investing protein in one of the three reactions of the model in Figure 1 implies that the growth rate is zero. We also know just from continuous dependence that the growth rate will reach a maximum somewhere in the interior of the domain in which *e*_1_ + *e*_2_ + *e*_*r*_ = *e*_*T*_, *e*_*i*_ ≥ 0. But here we show that the growth rate actually never has any local maxima: in fact, it is concave (as Figure 2 attests).

Three properties of our model—increasing enzyme concentrations improves the flux; increasing nutrient concentration does as well; optimizing growth rate can be formulated as a convex optimisation problem— essentially guarantee that the growth rate is a concave function of enzyme concentrations. This implies that a given suboptimal set of enzyme concentrations may always be slightly altered, without using more enzyme in total, in order to increase the growth rate. There is no risk of local maxima. This statement holds for very large kinetically explicit elementary metabolic networks, see Theorem 1.

#### Estimating optimal enzyme concentrations on the basis of current growth rate

Let us now consider a situation in which the enzyme concentrations are suboptimal, for instance because the nutrient concentration has suddenly changed. These concentrations give rise to the current, instantaneous growth rate *λ*(***e***; *<u>x</u>*_<u>0</u>_).

From the above arguments, there exists a set of optimal steady state enzyme concentrations with corresponding maximal steady state growth rate that is *equal to the current growth rate*: by decreasing the nutrient concentration sufficiently, the attainable maximal growth rate decreases until it reaches the current (suboptimal) growth rate, see Figure 3. This set of optimal enzyme concentrations is ***e***^*est*^(*λ*). The nutrient concentration at this optimizer is lower than the currently prevailing one if and only if the current enzyme concentrations are not optimal.

In short, there exists a unique *<u>ξ</u>*_<u>0</u>_ *< <u>x</u>*_<u>0</u>_ such that

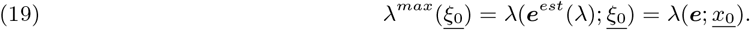

Second, by eq. (13) these estimated optimal concentrations induce a higher growth rate in the current nutrient condition than the current (suboptimal) enzyme concentrations do,

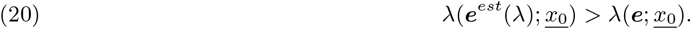

This is the property that will ensure that the growth rate acts as a Lyapunov function.

#### Characteristics of robust growth-rate optimizing gene networks

We now turn to dynamics. Recall eq. (1) and (4),

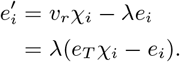

The ribosomal allocation parameters *χ*_*i*_ are assumed to be induced by a gene regulatory network, and are still to be defined: they are to be a function of the metabolic state of the cell.

Let us assume that the metabolic network is coupled to a gene network that is capable of sustaining a maximal steady state growth rate. It induces enzyme synthesis rates *v*_*r*_***χ*** such that at steady state the enzyme concentrations are optimal. Then if the nutrient condition is *<u>x</u>*_<u>0</u>_, at steady state we must have

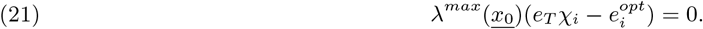

so that

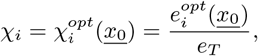

as derived in eq. (12).

We now come to the very heart of the paper: if the gene network inducing the ribosomal allocation by gene expression uses only the current growth rate as input to determine the enzyme synthesis rates, we have no choice but to set

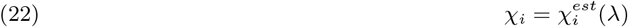

and thus enzyme concentrations change according to

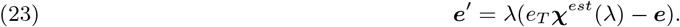

The reason why eq. (22) is the only option is that by eq. (18) it ensures that at steady state, the pathway is in its optimal growth state.

To summarise, the complete equations for the model in Figure 1 are given by

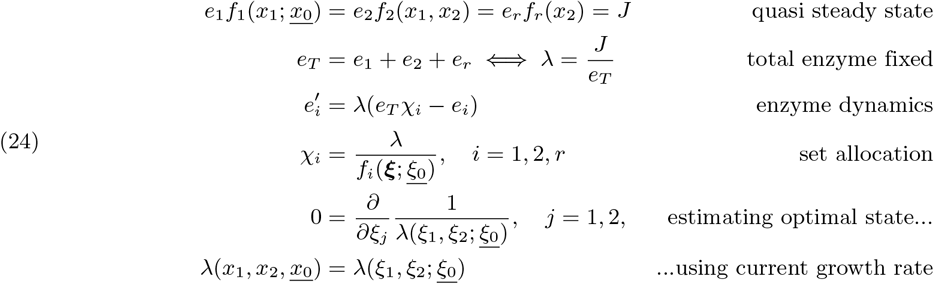

The first line sets *x*_1_, *x*_2_ and *J* as a function of ***e***. The last three lines are six equations, in six unknowns, *χ*_1_, *χ*_2_, *χ*_*r*_, *ξ*_0_, *ξ*_1_ and *ξ*_2_, treating *<u>x</u>*_<u>0</u>_, *x*_1_, *x*_2_ as parameters. In the SI we give two other formulations of the last two lines in system (24).

It is important to note that at this point it is not at all obvious that system (24) actually works. All that is clear so far is that if the dynamics reaches a steady state, this state is necessarily one of maximal growth—this is true by design.

So let us now turn to the actual dynamics of the metabolic-enzyme system with gene expression controlled according to system (24).

#### Stability and robustness of the adaptive control

The main open question is: how can we be sure that the dynamics of system (24) always result in ***e***(*t*) converging towards ***e***^*opt*^(*<u>x</u>*_<u>0</u>_)?

Using the relation ***χ***^*est*^ = ***e***^*est*^*/e*_*T*_, eq. (23) may be written as

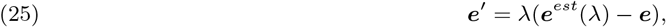

indicating that ***e*** moves directly towards ***e***^*est*^(*λ*) at any moment in time. Moreover, since the growth rate at ***e***^*est*^(*λ*) is higher than the current growth rate (see eq. (20), and Figures 2B, 3), the growth rate invariably increases: it is therefore a Lyapunov function. As long as the balance between synthesis and dilution in eq. (25) has not been reached, the enzyme concentrations keep changing, the growth rate increases, the corresponding enzyme synthesis rates changes, and this cycle keeps repeating. The complete formal statement is presented in Theorem 2.

The only steady state this coupled metabolite-enzyme system can reach is one in which the growth rate is actually the maximal one corresponding to the current environmental conditions, i.e., eq. (18) holds. The system is still robust to changes in these conditions, since the gene network still uses the instantaneous growth rate as input, see eq. (22).

## Dynamics of the model in Figure 1: robust optimisation of growth rate across conditions

For the linear metabolic network in Figure 1A an example simulation is shown in Figure 4.

For each nutrient condition *<u>x</u>*_<u>0</u>_, the optimal ribosomal allocation and corresponding maximal growth rates were computed and plotted against each other (Figure 4B). Note, for example, that to achieve a higher maximal growth rate, enzyme 1 should be expressed more with respect to enzyme 2, whilst ribosomal protein remains relatively constant (these relations depend on the kinetic constants in the rate laws).

Figure 4B specifies the action of the gene network, using current growth rate as input, and inducing enzyme synthesis rates according to the ribosomal allocation functions shown there. As shown in Theorem 2, for this gene network control, the growth rate is itself a Lyapunov function. After any change in the nutrient condition, the growth rate first drops to a new quasi steady state, but then always increases until a new maximum is reached (Figure 4C). Since the input-output relation (Figure 4B) does not depend on the nutrient concentration, the metabolic network is able to adapt after a change in nutrients and hones in on a new optimal steady state (Figure 4C, D), thus showing that this control is robust to nutrient availability.

In Theorem 2 we show that this robust adaptation towards maximal growth rate is not confined to the toy model illustrated here, but is in fact a pervasive property of elementary whole-cell metabolic networks.

### Cellular perception of the growth rate using the alarmone ppGpp

The growth rate of the cell is a systemic property, the result of the combined processes of metabolism, gene expression and protein synthesis. It is thus not a quantity cells can have direct access to. It is, however, possible that the cell tracks the growth rate via a metabolite concentration that is in a faithful one-to-one correspondence with it, even in dynamically changing conditions. Such a metabolite could then be used to control a gene regulatory network, as a proxy for the growth rate itself. As we now show, the way ppGpp is used by *E. coli* to control gene expression is to a large extent in agreement with the mechanism discussed in this paper.

ppGpp is a key signaling metabolite involved in the regulation of ribosome biosynthesis, and in coordinating responses to changes in environmental conditions [46]. It has been shown experimentally that the ppGpp concentration is directly coupled to the rate of translational elongation (and thus, to the protein synthesis rate *v*_*r*_), both in dynamic and constant conditions [62]. The relation between the ppGpp concentration *g*(*t*), measured relative to its concentration before a nutrient perturbation, and the inverse of elongation rate *ER*(*t*) is linear (Figure 5A, [62]). This linear relationship is given by

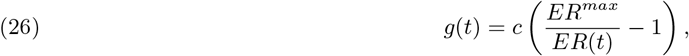

where *c* ≈ 4.0 and *ER*^*max*^ ≈ 19.4 *aa/s* are condition-independent constants. In balanced growth, this extends to a fixed relationship between growth rate and ppGpp (Figure 5B, [62]).

**Figure 5.**
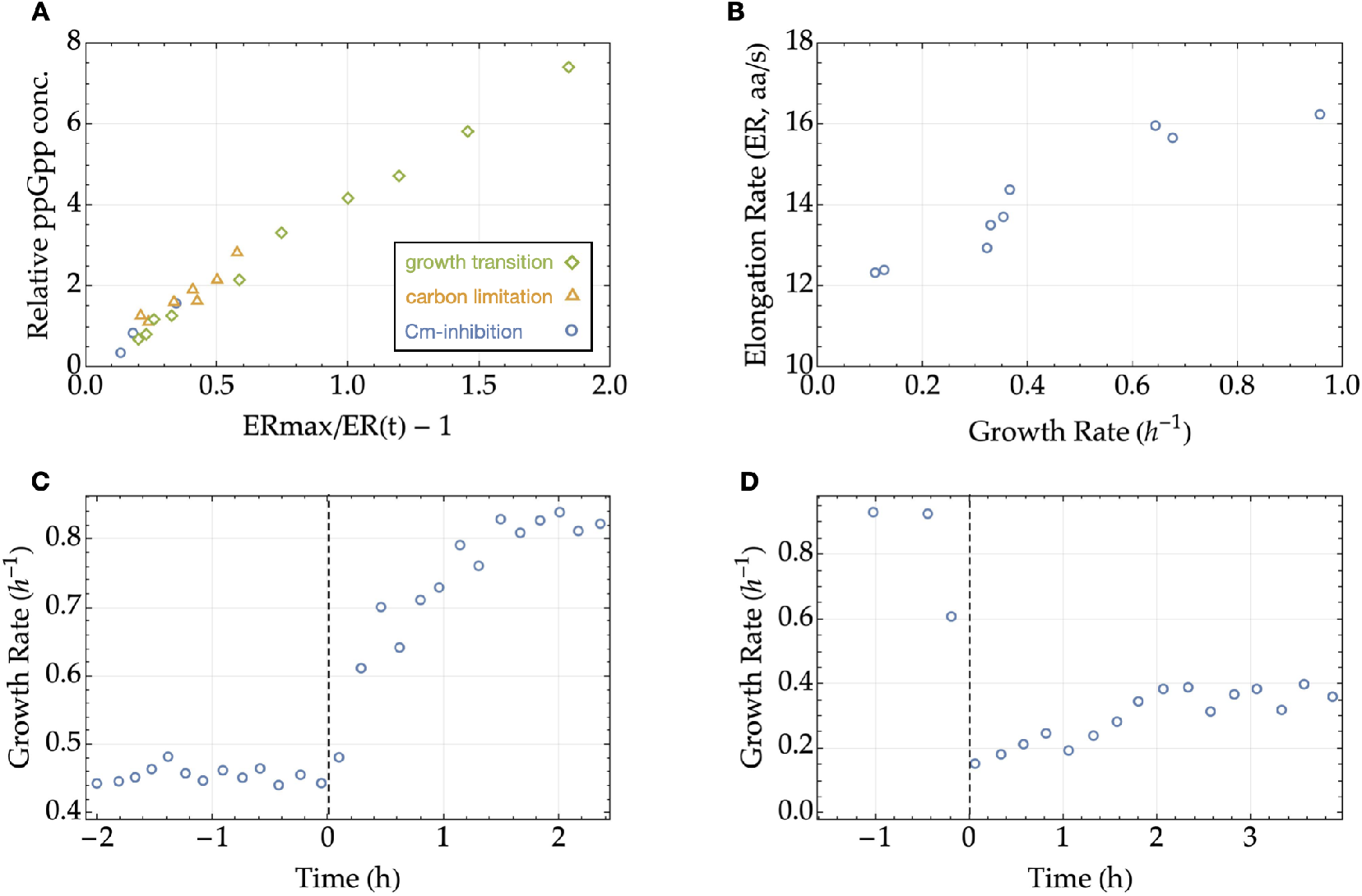
Experimental evidence supporting adaptive control. A: In steady state and dynamic conditions, inverse elongation rate *ER*(*t*) and relative ppGpp concentration are linearly related, approximately according to eq. (26) The relation holds during dynamic growth transitions, in carbon-limited growth and when translation is inhibited using sublethal doses of chloramphenicol (Cm). B: In balanced growth, the growth rate and elongation rate are positively related. C, D: Immediate increase of growth rate in *E. coli* after nutrient upshift (C) or downshift (D). At *t* = 0, a nutrient is added (left) or deleted (right) from the medium. Note that the growth rate increases directly after the change in shift, consistent with the idea proposed in this paper that the growth rate acts as a Lyapunov function, as illustrated in Fig. 4C. See [33, 37, 40, 44] for other experimental nutrient upshift examples, and [65, 40] for downshift examples, all showing the same behaviour. A and B adapted from [62] Fig 2B and D; C and D adapted from [18], Fig 1b and 1g. See those papers for experimental details.

In the parlance of this paper, ppGpp is thus used to control gene expression using a steady state estimator: the ribosomal allocation is directly determined by the current ppGpp concentration, which corresponds to a unique *steady state* growth rate. The dynamical change in enzyme concentrations only stops when then current growth rate is in fact equal to the growth rate corresponding to the current ppGpp concentration.

This idea is central to the dynamic model by Erickson et al. [18], and also to ours. The main difference between the two approaches is that in [18], the balanced growth relation between ppGpp, translation elongation rate and growth rate, was based on measurements of cultures in balanced growth, and did not involve any optimisation. In our case, the ribosomal allocation, as a function of the instantaneous growth rate (or a proxy thereof) is uniquely *derived* from the kinetics of the metabolic rate laws. Thus, we provide a mechanistic explanation while Erickson et al. [18] provided a phenomenological description.

As predicted by the theory presented here, after a nutrient shift the growth rate changes quickly to a new quasi steady state, after which a gradual increase in the growth rate is observed. See Figure 5C, D for an example of both a nutrient upshift and downshift. Other experimental examples may be found in [33, 37, 40, 44] for nutrient upshifts, and [65, 40] for downshifts. These data are thus consistent with the idea that the growth rate acts as a Lyapunov function, invariably increasing in time.

### Growth rate is concave in enzyme concentrations

We finish by stating the results more precisely, using the notation for general metabolic networks given in the Methods. Proofs may be found in the SI. The first theorem shows that for many elementary metabolic networks the specific flux is concave in enzyme concentrations.

#### Theorem 1.

*Consider an elementary pathway with n enzymatic reactions and one ribosomal reaction, with steady state reaction rates v*_*i*_ = *JV*_*i*_ = *e*_*i*_*f*_*i*_(***x***, *<u>x</u>*_<u>0</u>_), *i* = 1, …, *n, r and with <u>x</u>*_<u>0</u>_ *a fixed nutrient concentration. Let J be the protein synthesis flux through the pathway for some enzyme concentration vector, and let λ* = *J/e*_*T*_ *be the cellular growth rate. Assume that*

1. *For each enzyme allocation* ***e*** *and nutrient concentration <u>x</u>*_<u>0</u>_, *there exists a unique quasi steady state of metabolite concentrations* ***x***;
2. *an increase in <u>x</u>*_<u>0</u>_ *results in an increase in J, at fixed enzyme concentrations* ***e***;
3. *an increase in any enzyme concentration e*_*k*_ *results in an increase of J, at fixed nutrient concentration <u>x</u>*_<u>0</u>_*(and keeping all other e*_*j*_ *fixed);*
4. *the function*

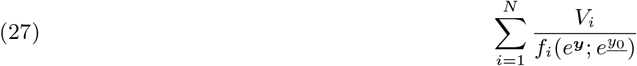

*is (strictly) convex in* ***y*** *on the domain in which all f*_*i*_ *functions are positive*. *Then λ* = *J/e*_*T*_ *is (strictly) concave in* ***e***.

Assumptions 2 and 3 are general, intuitive properties of metabolic networks: more nutrient yields a higher flux, and so does investing more enzyme in any one reaction. The function in Assumption 4 is the reciprocal of the growth rate (cf. eq. (7)) at quasi steady state in logarithmic variables. It has been shown to be strictly convex in elementary networks for a very large class of rate laws [41, 45].

### Adaptively controlled elementary networks always converge to the optimal steady state

The following theorem builds on the previous one, and contains the main result of this paper.

#### Theorem 2.

*Consider an elementary metabolic pathway with n enzymatic and one ribosomal reactions and steady state reaction rates v*_*i*_ = *JV*_*i*_ = *e*_*i*_*f*_*i*_(***x***, *<u>x</u>*_<u>0</u>_), *i* = 1 …, *n, r with <u>x</u>*_<u>0</u>_ *a nutrient concentration. Assume that enzymes and ribosome change according to system (24). Assume furthermore that*

1. *λ* = *J/e*_*T*_ *is concave in* ***e***,
2. *An increase in <u>x</u>*_<u>0</u>_ *induces a higher flux at fixed enzyme concentrations* ***e***. *Then λ is a Lyapunov function, and increases along all orbits. Moreover, the dynamical system is globally stable on*

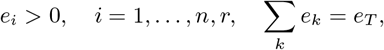

(***x***(*t*), ***e***(*t*)) → (***x***^*opt*^, ***e***^*opt*^), *with the optimum defined by <u>x</u>*_<u>0</u>_, *and λ*(*t*) ↗ *λ*^*max*^(*<u>x</u>*_<u>0</u>_).

Theorems 1 and 2 together show that an elementary metabolic network whose enzyme concentrations are determined by the proposed adaptive control, using growth rate as input, attains the maximal steady state growth rate robustly across conditions.

## Discussion

Cells display dynamical responses of protein expression to remain competitive across changing environmental conditions. In this paper we have shown that a gene regulatory feedback that is based on a readout of the cellular growth rate provides a stable and robust way to implement such a response. The stability of the response is ensured because the growth rate acts as a Lyapunov function and always increases after a nutrient perturbation. *E. coli* seems to closely approximate this mechanism using ppGpp as a proxy for the instantaneous protein translation rate.

### Optimality of protein expression

There is an ongoing debate to what extent cells optimise protein expression. Cells experimentally often seem to tune protein concentration to maximal growth rate [28, 32, 30, 57, 48], and these regulation strategies are hard wired [1]. At low nutrient concentrations (and thus low growth rates), however, the fraction of ribosomes that are actively translating new protein is kept low by active sequestration [37, 62] (this does not preclude optimisation, as the active ribosomes may still be optimally allocated [6]); moreover, protein expression is diverted to preparing for adverse conditions and nutrient changes [6].

### Modelling approaches

Adaptation to changing conditions by microorganisms through dynamic reallocation of biosynthetic resources has grown into a rich field the last few years, with different approaches and viewpoints.

Using experimentally measured protein allocation from balanced growth cultures, excellent quantitative agreement in dynamic transitions were shown to result when applying those relations in a theoretical feedback control based on ppGpp regulation [18]. This approach does not use optimisation as the underlying assumption, but the logic behind the control is identical to the one presented here, with steady state relations being used as the control law for dynamic conditions. Instead of deriving the input-output relations from optimality principles, as done here, they were measured. The ppGpp-based feedback has recently been further characterised experimentally [62], which resulted in an improved dynamic theoretical framework [15].

Optimality has been incorporated in different ways. Several studies, including the present one, focus on dynamics towards maximal balanced growth rate (or more generally, flux per unit enzyme) [3, 33, 45, 8]. This is a form of adaptive control. Some have focused on dynamic regulation for a fixed condition (so towards a fixed optimal state set by the environmental condition) [33, 8], while others have also considered robustness to changes in those conditions [3, 45]. There is also a growing literature focusing on optimal control [43, 59, 22, 27, 63]. Here, the objective is to maximise biomass accumulation within a fixed time period using dynamic regulation, rather than achieving a maximal balanced growth rate.

The idea of flux sensing was studied experimentally in *E. coli* [34, 31]. This work does not involve optimality, but provided impetus to modelling of dynamical regulation using metabolite-binding transcription factors, first in galactose uptake in *S. cerevisiae* [3] and then in more general metabolic networks [45].

### Elementary Flux Modes

We have explained the theory using a coarse grained model (Figure 1), and discussed its relevance for control of the balance between metabolism and protein synthesis with the alarmone ppGpp in *E. coli*. Coarse grained models based on growth laws [51, 54, 2] are currently the norm in the field [4, 56, 8, 15, 18, 33]. These models have been very insightful, but greatly simplify the details of larger metabolic networks. They all have the property that each and every reaction is required to sustain a flux. One flux value then determines all flux values: they are one-degree-of-freedom pathways. Such minimal networks are called Elementary Flux Modes [52, 53, 21], and they appear naturally in the context of large (genome-scale) metabolic networks as growth rate maximisers [61, 38, 11]. The theory presented in this paper applies to all such elementary networks, with quite arbitrary reaction kinetics [41, 45]. It thus also provides insight into, for instance, catabolic uptake pathways, as the detailed kinetic model of galactose uptake in yeast exemplifies [3].

In this paper we have assumed that metabolism settles down to a quasi steady state at a rate that is much higher than the rate of change of protein synthesis. This idealisation is currently necessary to prove stability of the controlled pathway, but not for the construction of the control. Indeed, in [45] we have presented a more general theory in which the metabolic QSS assumption is dropped, and in which the gene network uses metabolite-binding transcription factors as input to control protein synthesis. Simulations suggest that this type of control is equally robust, but a mathematical proof is currently beyond our reach. The growth rate is then definitely not a Lyapunov function, however, and stable oscillations may appear, as also reported by Droghetti et al. [15].

### ppGpp regulation

This paper provides a new view on why ppGpp directly measures the ribosomal translation rate of cells. We suggest that since the translation rate is proportional to growth rate [62], its measurement by ppGpp makes the growth rate of a cell a Lyapunov function for the combined system of catabolism, anabolism and the regulation of their genes. We already knew that ppGpp coordinates the balance between catabolic and anabolic protein expression by allocating RNAP-sigma 70 over the associated operons, and even co-regulating the balance between growth-associated and growth-unassociated functions of the cells (e.g. stress tolerance and environmental sensing) [46]. From the perspective of this paper, ppGpp can be viewed as a form of feedback regulation using the current performance (the growth rate) to control the dynamic system responsible for this performance (metabolism) by setting the balance between metabolic and ribosomal protein synthesis. Moreover, we show that this construction ensures stability of the dynamic response: microbes remain competitive even when faced with changing nutrient conditions that require adaptive responses. That something as complex and multifaceted as growth rate is compressible into a single variable is an example of dimension reduction that perhaps make systems as complex as cells controllable [62]. Here we provide further evidence of its benefits.

### Lyapunov functions in control theory

To stabilise complex nonlinear systems such as an aeroplane, engineers often design controllers in such a way that a suitable function can be shown to be a Lyapunov function for the controlled system [23, e.g.]. Lyapunov functions only decrease or increase in value over time, along any orbit of the controlled dynamical system. The existence of such a Lyapunov function thus ensures the stability of the controlled system, making it a powerful technique. However, there is no general recipe for finding them. In the biological case considered here, we are in the unusual situation that the input to the controller (the cellular growth rate) always increases over time, and is thus itself a Lyapunov function.

With this paper, we have given compelling evidence that even living cells might be able to exploit the stabilising control mechanism of Lyapunov functions by evolutionary moulding and improved adaptation to changing conditions.

## Methods

We focus on whole-cell models that lead to cellular growth. The synthesis of enzymes and ribosomes from precursors (amino acids, etc.) is thus included. We neglect proteins involved in maintenance tasks (the so-called Q-sector [56, 35, 63, e.g.], which is not regulated by ppGpp).

We model the change in metabolite concentrations ***x*** in the metabolic network by

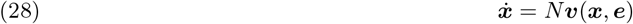

where *N* is the metabolic stoichiometric matrix, ***v*** the vector of reaction rates, and ***e*** the vector of enzyme concentrations. As the last reaction in metabolism we invariably choose the protein translation reaction by the ribosome, and indicate it with subscript *r*: *v*_*r*_. Metabolism generally quickly settles into a quasi steady state; this we assume throughout: *N* ***v***(***x***; ***e***) = **0**.

We model the rate *v*_*i*_ through reaction *i* by [10]

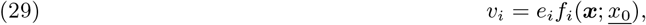

where *e*_*i*_ is the concentration of enzyme *i* and *f*_*i*_(***x***; *<u>x</u>*_<u>0</u>_) incorporates the nonlinear dependence on metabolite concentrations ***x*** and an external nutrient concentration *<u>x</u>*_<u>0</u>_ assumed to be fixed. (Of course, each *f*_*i*_ generally depends on only a few *x*_*j*_ and maybe one or two on *<u>x</u>*_<u>0</u>_.)

Elementary Flux Modes (EFMs) are subnetworks that are made by deleting nonparticipating metabolites and reactions from eq. (28) and reducing the stoichiometric matrix accordingly. This is to be done such that all remaining reactions are essential to sustain a steady state flux *J* [52, 53, 21]; deleting any further reaction by not producing the corresponding enzyme stops the flux through the pathway. EFMs have “one flux-degree-of-freedom” [58, 29]: prescribing one flux value immediately determines all flux values. In other words, *N* ***v*** = **0** is then replaced by ***v*** = *J* ***V***, where ***V*** = (*V*_1_, …, *V*_*n*_, *V*_*r*_) is a fixed vector determined by the stoichiometry of the EFM. (Concrete examples may be found in [5].) It is uniquely defined by choosing *V*_*r*_ = 1, without loss of generality. The steady state flux *J* is determined by solving, for a given set of enzyme concentrations (*e*_1_, …, *e*_*n*_), an additional ribosome concentration *e*_*r*_ and external nutrient concentration *<u>x</u>*_<u>0</u>_,

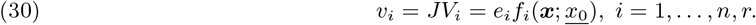

The set of possible flux vectors ***v*** = *J* ***V*** thus forms a fixed straight line through the origin, with a direction determined by ***V***. The line is parameterized by *J*. Since in this paper we focus on one particular EFM, we may choose ***V*** to be a strictly positive vector. For a positive flux, all kinetics functions *f*_1_, …, *f*_*n*_, *f*_*r*_ must therefore also be positive.

By eq. (29) all reaction rates scale linearly with enzyme concentration. As a result, *J* is 1-homogeneous in ***e*** [24]: *J* (*r****e***) = *rJ* (***e***) for any *r >* 0. With *e*_*T*_ denoting the total cellular protein concentration, the specific flux *J* (***e***)*/e*_*T*_ is thus 0-homogeneous in ***e***: *J* (*r****e***)*/*(*re*_*T*_) = *J* (***e***)*/e*_*T*_.

Cells grow in volume as new cellular material is being produced. This causes dilution of cellular concentrations. For metabolite concentrations, dilution by growth is generally not taken into account because metabolic fluxes are orders of magnitude larger [13]. We take the same approach here. For enzyme concentrations, however, enzyme synthesis rates and dilution rates are comparable. We therefore assume that enzyme concentrations change according to

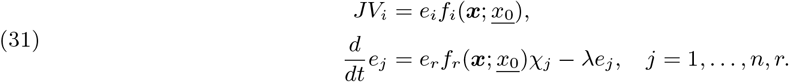

Here, *e*_*r*_ is the ribosome concentration, *f*_*r*_(***x***; *<u>x</u>*_<u>0</u>_) the saturation level of ribosomes (including the catalytic rate constant), and *χ*_*j*_ is the fraction of active ribosomes transcribing enzyme *j*, or the ribosome if *j* = *r*. The dependence of *f*_*r*_ on *<u>x</u>*_<u>0</u>_ is for notational consistency, and to highlight that the performance of the metabolic network still depends on the nutrient concentration *<u>x</u>*_<u>0</u>_. The growth rate is denoted by *λ*. We assume that the *χ*_*j*_ sum to one, and that the total protein content is constant,

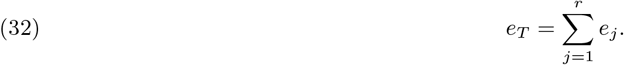

Then we see that [13, 11, 14]

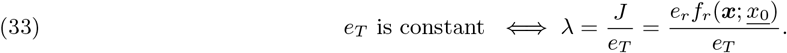

So the cellular growth rate is the total protein synthesis rate (which is the flux *J*, since *V*_*r*_ = 1) divided by the total protein concentration, a specific flux.

In this paper, we use the word *allocation* to specify a vector up to a scaling. The vectors ***e*** and 2***e*** thus indicate the same allocation of enzymes to a metabolic network. Since specific flux is 0-homogeneous in ***e***, it is determined by the enzyme allocation.

## Supporting information

SI Matlab and Mathematica files

## Acknowledgments

The authors thank Gosse Overal for his work on early attempts to prove global stability. We also thank Marco Cosentino Lagomarsino, Daan de Groot, Terry Hwa, Wolfram Liebermeister, Elad Noor, Ralf Steuer, Bas Teusink, and Pieter Rein ten Wolde for many insightful discussions.

