## Supplementary Text for "Sensing cellular growth rate facilitates its robust optimal adaptation to changing conditions"

### SUPPLEMENTARY INFORMATION FOR 'SENSING CELLULAR GROWTH RATE FACILITATES ITS ROBUST OPTIMAL ADAPTATION TO CHANGING CONDITIONS'

ROBERT PLANQUÉ[1,*], JOSEPHUS HULSHOF[1], AND FRANK J. BRUGGEMAN[2]

#### 1. Concavity of specific flux

##### 1.1. Why would $J(e)/e_T$ be concave? The relationship with the harmonic mean.

In this section, we focus only on the specific flux through an Elementary Flux Mode, without coupling it to enzyme dynamics and qORAC control. We therefore do not need to distinguish between enzymes and the ribosome: the ribosome is just last enzymatic reaction in the EFM. So in this section, an EFM has $n$ reactions (rather than $n+1$ as in the main text), each catalyzed by an enzyme with concentration $e_1, \ldots, e_n$.

Elementary Flux Modes have "one flux-degree-of-freedom". The steady state flux is determined by solving

$$v_i = JV_i = e_i f_i(\boldsymbol{x}). \tag{1}$$

Here the ratios $V_1 : V_2 : \cdots : V_n$ are fixed and determined by the stoichiometry of the EFM. Because we focus on one EFM, we may choose $\boldsymbol{V}$ to be a strictly positive vector. To fix $J$, we choose $V_n = 1$ without loss of generality. The set of possible flux vectors form a fixed straight line through the origin, with a direction determined by $\boldsymbol{V}$. The line is parameterized by $J$.

A naive approach to showing that $J(\boldsymbol{e})/e_T$ is concave would entail to write $\boldsymbol{x} = \boldsymbol{x}(\boldsymbol{e})$ and $J = J(\boldsymbol{e})$ in eq. (1), and perform implicit differentiation to second order to construct the Hessian $H_J(\boldsymbol{e})$. This works only for very small metabolic pathways; for the amusement of the reader, we have supplied an example for the linear chain in the Section 1.6. It should be immediately obvious why this line of proof cannot be generalized to general EFMs with arbitrary kinetic rate laws.

Writing eq. (1) first as

$$\frac{e_i}{J} = \frac{V_i}{f_i(\boldsymbol{x})}, \tag{2}$$

and summing over all $i = 1, \ldots, n$ we find

$$\frac{e_1 + \cdots + e_n}{J} = \sum_i \frac{V_i}{f_i(\boldsymbol{x})},$$

so that for the specific flux

$$\frac{J}{e_1 + \cdots + e_n} = \frac{1}{\sum_i \frac{V_i}{f_i(\boldsymbol{x})}}. \tag{3}$$

Introducing notation for the harmonic mean of $\boldsymbol{x}$ by

$$H(x_1, \ldots, x_n) \equiv \frac{n}{\frac{1}{x_1} + \cdots + \frac{1}{x_n}} = \frac{1}{\left\langle \frac{1}{\boldsymbol{x}} \right\rangle},$$

we may write eq. (3) as

$$J = \langle \boldsymbol{e} \rangle \, H(\boldsymbol{f}/\boldsymbol{V}).$$

(Here, the division $\boldsymbol{f}/\boldsymbol{V}$ should be interpreted elementwise.) The function

$$O(\boldsymbol{x}) \equiv \sum_i \frac{V_i}{f_i(\boldsymbol{x})}, \tag{4}$$

sometimes referred to as the enzyme cost function from Enzyme Cost Minimization [2] or simply the objective function [4], plays an important part in our current understanding of the optimization problem of maximal specific flux in EFMs. In logarithmic coordinates, $\boldsymbol{y} = \ln \boldsymbol{x}$, this function has been shown to be strictly convex on a convex bounded domain for a wide variety of rate laws [2, 4]. This shows that optimizing specific flux (using the metabolite concentrations as optimization variables) is a well-posed problem and has a unique solution, but it does not offer much insight into the dependence of $J$ on $\boldsymbol{e}$.

A new view on $J$ as a function of $\boldsymbol{e}$ may be obtained by observing the symmetry between $e_i$ and $f_i$ in eq. (1). We can of course also rewrite eq. (1) to

$$\frac{f_i(\boldsymbol{x})}{JV_i} = \frac{1}{e_i},$$

to obtain (after summing and taking reciprocals again)

$$J = \langle \boldsymbol{f}(\boldsymbol{x})/\boldsymbol{V} \rangle \, H(\boldsymbol{e}). \tag{5}$$

Of course the steady state metabolite concentrations $\boldsymbol{x}$ are determined by the allocation of enzymes across the pathway, i.e., $\boldsymbol{x} = \boldsymbol{x}(\boldsymbol{e})$. The above expression therefore does not provide us with an explicit expression for $J$ as a function of $\boldsymbol{e}$. But note that the harmonic mean is a concave function. It is not strictly concave in all directions: it is also 1-homogeneous. But in all directions other than along lines through the origin, it is strictly concave. In particular, it is strictly concave on the enzyme simplex $e_1 + \cdots + e_n = e_T$. Therefore, on this simplex also the specific flux $J(\boldsymbol{e})/e_T$ is strictly concave.

So in conclusion, even though eq. (5) only defines $J$ implicitly as a function of $\boldsymbol{e}$, we do obtain an intuition why $J(\boldsymbol{e})/e_T$ could indeed be strictly concave: if the mean (scaled) rate functions do not vary much while changing $\boldsymbol{e}$, then indeed $J(\boldsymbol{e})/e_T$ is strictly concave.

1.2. **The metabolic steady state map.** The foregoing discussion is completely general, and since $\boldsymbol{x} = \boldsymbol{x}(\boldsymbol{e})$ in quasi steady state, this functional relationship is usually a purely implicit one. We have therefore only gained a crude intuition so far. In a simple example, however, we can make it explicit, and also reveal that concavity of $J$ might be related to convexity of a metabolite steady state map.

Consider

$$\underline{x_0} \overset{e_1}{\rightleftharpoons} x_1 \overset{e_2}{\rightleftharpoons} x_2 \overset{e_3}{\rightleftharpoons} x_3 \overset{e_4}{\rightleftharpoons} \cdots \overset{e_{n-1}}{\rightleftharpoons} x_{n-1} \overset{e_n}{\rightleftharpoons} \underline{x_n}.$$

If we assume linear kinetics $f_i = x_{i-1} - x_i$, where $x_0 = \underline{x_0}$ and $x_n = \underline{x_n}$ are assumed to be fixed, then eq. (5) becomes

$$J = (\underline{x_0} - \underline{x_n}) \frac{1}{\frac{1}{e_1} + \cdots + \frac{1}{e_n}}. \tag{6}$$

Strict concavity of $J(\boldsymbol{e})/e_T$ can be concluded directly.

We can now define a metabolite steady state map in the following way. Fixing again $x_n = \underline{x_n}$, the last steady state equation is

$$e_n(x_{n-1} - \underline{x_n}) = J.$$

Introducing $z_i = e_i/J$, this reads $z_n(x_{n-1} - \underline{x_n}) = 1$, and we see that the steady state concentration $x_{n-1}$ satisfies

$$x_{n-1} = \frac{1}{z_n} + \underline{x_n}.$$

By going back progressively towards the start of the chain of reactions, we find

$$\begin{aligned} x_{n-2} &= \frac{1}{z_{n-1}} + x_{n-1} \\ &= \frac{1}{z_{n-1}} + \frac{1}{z_n} + \underline{x_n} \end{aligned}$$

and so on, so that

$$\underline{x_0} - \underline{x_n} = F(\boldsymbol{z}) \equiv \frac{1}{z_1} + \frac{1}{z_2} + \cdots + \frac{1}{z_n}. \tag{7}$$

This is, of course, nothing but a different way of writing eq. (6). The metabolite steady state map $F(\boldsymbol{z})$ is essentially the reciprocal of the harmonic mean of $\boldsymbol{e}$.

Clearly, $F$ is a strictly decreasing and strictly convex function for $z_i > 0$. Next to that we have the objective function eq. (4)

$$O(\boldsymbol{x}; \underline{x_0}) = \frac{1}{\underline{x_0} - x_1} + \frac{1}{x_1 - x_2} + \cdots + \frac{1}{x_{n-1} - \underline{x_n}}, \tag{8}$$

which is strictly convex in logarithmic variables.

In the next section we generalize these ideas, and show that the convexity of eq. (8) is (largely) equivalent to convexity of eq. (7), and which in turn implies concavity of $J/e_T$.

1.3. **Existence and convexity of the metabolite steady state map.** The central concept we introduce to study both the concavity of specific flux and the adaptive control is the metabolite steady state map $F(\boldsymbol{z})$. First, recall that we set $z_i = e_i/J$. We wish to find, or at least define, a function $F(\boldsymbol{z})$ such that $\underline{x_0} = F(\boldsymbol{z})$ corresponds to the EFM being in quasi steady state.

1.3.1. *Existence.* For an arbitrary EFM with associated kinetic rate laws, we are not likely to be able to find the metabolite steady state map $F(\boldsymbol{z})$ explicitly. The steady state equations eq. (2) in $\boldsymbol{x}$ and $\boldsymbol{z}$ variables are

$$z_i f_i(x_1, \ldots, x_{n-1}; \underline{x_0}) = V_i, \quad i = 1, \ldots, n, \tag{9}$$

which is equivalent to

$$z_i = \frac{V_i}{f_i(x_1, \ldots, x_{n-1}; \underline{x_0})}. \tag{10}$$

Finding $F(\boldsymbol{z})$ amounts to inverting these maps, and finding functions $F_i(\boldsymbol{z})$ such that

$$x_i = F_i(\boldsymbol{z}), \quad i = 0, \ldots, n-1.$$

This may be done if the Inverse Function Theorem may be applied globally. Differentiating eq. (9) with respect to $z_j$, we find

$$0 = \delta_{ij} f_i(\boldsymbol{x}) + z_i \sum_{k=0}^{n-1} \frac{\partial f_i}{\partial x_k}(x_1(\boldsymbol{z}), \ldots, x_{n-1}(\boldsymbol{z}); \underline{x_0}(\boldsymbol{z})) \frac{\partial x_k}{\partial z_j}.$$

We thus need that the Jacobian $(\frac{\partial f_i}{\partial x_k})$ is invertible in each $\boldsymbol{x} = (x_1, \ldots, x_{n-1}; \underline{x_0})$.

The steady state map $F(\boldsymbol{z})$ can now be related to a different objective function, which has been known to have good convexity properties.

1.3.2. *Equivalent optimization problems.* To aid the reader, we start by summarising all the different optimization problems for maximal specific flux in an EFM.

Recall that the EFM pathway has stoichiometry matrix $N$, a set of positive flux ratios $V_1 : V_2 : \cdots : V_n$ where by convention we choose $V_n = 1$, and fixed rate laws $f_i(\boldsymbol{x}; \underline{x_0})$, one for each reaction. We also assume that the metabolite steady state map $\underline{x_0} = F(\boldsymbol{z})$ exists. The fixed concentration $\underline{x_0}$ should be such that $J > 0$.

In the optimization problems that follow, we need to specify constraints, including the relevant domains for the optimization variables. The exact relevant domains may depend on the choice of EFM, the substrates and products involved in the different reactions, the existence of moieties, and the number of external concentrations that are fixed. We do not go into all these details here, for which see [2].

Reaction $i$ is assumed to involve substrate concentrations $\boldsymbol{x}_i^s$ and product concentrations $\boldsymbol{x}_i^p$. The inequality $\boldsymbol{x}_i^s \geq \boldsymbol{x}_i^p$ should be interpreted loosely. E.g., for a reaction $2A \to B$, the numerator of the rate law would contain a term $a^2 - b/K_{eq}$, so for positive flux through this reaction we would need to assume $a^2 > b/K_{eq}$.

We now have the following equivalent optimization problems for this EFM. The central one is eq. (14), but to elucidate how to rewrite it into the other forms, we have put it in the middle rather than at the start.

$$\min_{\boldsymbol{z}}\left\{\sum_i z_i \;\middle|\; F(\boldsymbol{z}) = \underline{x_0}\right\} \tag{11}$$

$$\Longleftrightarrow$$

$$\min_{\boldsymbol{e},J}\left\{\frac{\sum_i e_i}{J} \;\middle|\; F\left(\frac{\boldsymbol{e}}{J}\right) = \underline{x_0},\ \boldsymbol{e} \geq \boldsymbol{0},\ J > 0\right\} \tag{12}$$

$$\Longleftrightarrow$$

$$\max_{\boldsymbol{e},J}\left\{\frac{J}{\sum_i e_i} \;\middle|\; F\left(\frac{\boldsymbol{e}}{J}\right) = \underline{x_0},\ \boldsymbol{e} \geq \boldsymbol{0},\ J > 0\right\} \tag{13}$$

$$\Longleftrightarrow$$

$$\max_{\boldsymbol{x},\boldsymbol{e}}\left\{\frac{J}{\sum_i e_i} \;\middle|\; N\boldsymbol{v} = \boldsymbol{0},\ v_i = e_i f_i(\boldsymbol{x}; \underline{x_0}),\ v_i \geq 0\right\} \tag{14}$$

$$\Longleftrightarrow$$

$$\min_{\boldsymbol{x},\boldsymbol{e}}\left\{\frac{\sum_i e_i}{J} \;\middle|\; v_i = JV_i,\ v_i = e_i f_i(\boldsymbol{x}; \underline{x_0}),\ v_i \geq 0\right\} \tag{15}$$

$$\Longleftrightarrow$$

$$\min_{\boldsymbol{x}}\left\{\sum_i \frac{JV_i/f_i(\boldsymbol{x}; \underline{x_0})}{J} \;\middle|\; \boldsymbol{x}_i^s \geq \boldsymbol{x}_i^p\right\} \tag{16}$$

$$\Longleftrightarrow$$

$$\min_{\boldsymbol{x}}\left\{\sum_i \frac{V_i}{f_i(\boldsymbol{x}; \underline{x_0})} \;\middle|\; \boldsymbol{x}_i^s \geq \boldsymbol{x}_i^p\right\} \tag{17}$$

$$\Longleftrightarrow$$

$$\min_{\boldsymbol{y}}\left\{\sum_i \frac{V_i}{f_i(e^{\boldsymbol{y}}; \underline{x_0})} \;\middle|\; \boldsymbol{y}_i^s \geq \boldsymbol{y}_i^p\right\}. \tag{18}$$

In this paper we have introduced eq. (11), whereas previously eq. (18) has been the focus of attention [2, 4]. Note that eq. (14) involves two sets of variables over which we should maximise, while eq. (11), eq. (17) and eq. (18) contain just one.

We could have added the following problems to the list above, by the homogeneity of $J(\boldsymbol{e})$. In eq. (15) we could choose $J = 1$,

$$\min_{\boldsymbol{x},\boldsymbol{e}}\left\{\sum_i e_i \;\middle|\; v_i = V_i,\ v_i = e_i f_i(\boldsymbol{x}; \underline{x_0}),\ v_i \geq 0\right\}, \tag{19}$$

and, equivalently, we could have chosen $e_T = 1$ and maximise $J$,

$$\max_{\boldsymbol{x},\boldsymbol{e}}\left\{J \;\middle|\; v_i = JV_i,\ v_i = e_i f_i(\boldsymbol{x}; \underline{x_0}),\ \sum_i e_i = 1,\ v_i \geq 0\right\}. \tag{20}$$

1.3.3. *(Strict) convexity/concavity of the metabolite steady state map.* In [2] and [4], it was shown that the objective function in eq. (18) is strictly convex for a large class of rate laws. There is a direct connection between metabolite steady state map $F(\boldsymbol{z})$ and the old objective function. From eq. (10), note that

$$\sum_{i=1}^{n} z_i = \sum_{i=1}^{n} \frac{V_i}{f_i(e^{y_1}, \ldots, e^{y_{n-1}}; e^{y_0})}$$

is convex in $\boldsymbol{y}$. Indeed, each of the components

$$z_i = \frac{V_i}{f_i(e^{y_1}, \ldots, e^{y_{n-1}}; e^{y_0})}$$

is convex in the relevant $y_j$ variables. We will show: if each component of $\boldsymbol{z} = h(\boldsymbol{y})$ is convex and $h$ is invertible, then each component function of

$$(x_0, \ldots, x_{n-1}) = (e^{y_0}, \ldots, e^{y_{n-1}}) = (e^{h_1^{-1}(\boldsymbol{z})}, \ldots, e^{h_n^{-1}(\boldsymbol{z})})$$

is convex. We actually only need that the first component $x_0 = \exp h_1^{-1}(\boldsymbol{z}) = F(\boldsymbol{z})$ is convex.

In [3] it was shown that the inverse of an invertible convex function $f : D \subseteq \mathbb{R}^n \to R \subseteq \mathbb{R}^n$ is itself convex when each component of the inverse $g$ is decreasing in all of its variables (i.e., if $\nabla g(\boldsymbol{y}) \in \mathbb{R}^n_{<0}$). Indeed, for any component function $g_m$ we have that

$$Df^T H_{g_m} Df$$

is positive definite, so that the Hessian $H_{g_m}$ is also positive definite [3] (if $B$ is invertible then $A$ is positive definite if and only if $B^T AB$ is).

We can further deduce that if $g_m$ is convex, then so is $e^{g_m}$: taking derivatives of $e^{g_m(\boldsymbol{y})}$ in the direction of $y_p$ and then to $y_q$, $p, q = 1, \ldots, N$ yields

$$e^{g_m(\boldsymbol{y})} \left( H_{g_m}(\boldsymbol{y}) + \nabla g_m(\boldsymbol{y}) \nabla g_m^T(\boldsymbol{y}) \right). \tag{21}$$

Since $f$ and $g$ are each other's inverses, the Jacobian matrices are each other's inverses as well,

$$Df Dg = I,$$

so that for one component we have

$$\nabla g_m^T Df = \boldsymbol{e}_m^T$$

(where with a bit of abuse of notation $\boldsymbol{e}_m$ is now the $m$-th standard basis vector filled with zeros and one 1 on the $m$-th position). Hence

$$Df^T \nabla g_m \nabla g_m^T Df = \boldsymbol{e}_m \boldsymbol{e}_m^T,$$

a matrix with one 1 in position $(m, m)$ and zeros elsewhere. Hence

$$Df^T \left( H_{g_m} + \nabla g_m \nabla g_m^T \right) Df$$

is still positive definite, and so is $H_{g_m} + \nabla g_m \nabla g_m^T$.

So we conclude that each $e^{h_i^{-1}}(\boldsymbol{z})$ is convex if it has a negative gradient everywhere. In particular, $F(\boldsymbol{z}) = e^{h_1^{-1}}(\boldsymbol{z})$ is convex when $\nabla F$ is everywhere negative.

**Convexity of $F(z)$ vs concavity of $J(e)/e_T$.** We now consider the general problem of relating convexity of $F$ to concavity of $J(\boldsymbol{e})/e_T$. Since the result is completely independent of the context, we use abstract notation here.

Consider the implicit function $\psi = \psi(\boldsymbol{x})$ defined by

$$F\left(\frac{\boldsymbol{x}}{\psi}\right) = 1.$$

Setting $z_i = x_i/\psi$, we wish to relate the Hessian of $F$ to the Hessian of $\psi$.

We use the following notation. $DF = DF(\boldsymbol{z})$ acts on $\boldsymbol{\xi}$ as

$$\langle DF, \boldsymbol{\xi} \rangle = \langle DF(\boldsymbol{z}), \boldsymbol{\xi} \rangle = F_k \xi_k = \nabla F \cdot \boldsymbol{\xi} = F_{\boldsymbol{\xi}}, \tag{22}$$

in which $F_k$ is the partial derivative of $F$ with respect to its $k$-th argument, summation convention of repeated indices is used, and $F_\xi$ is the directional derivative

$$F_{\boldsymbol{\xi}}(\boldsymbol{z}) = \lim_{t \to 0} \frac{F(\boldsymbol{z} + t\boldsymbol{\xi}) - F(\boldsymbol{z})}{t}$$

in $\boldsymbol{z}$.

Likewise

$$\langle D\psi, \boldsymbol{\xi} \rangle = \langle D\psi(x), \boldsymbol{\xi} \rangle = \psi_k \xi_k = \nabla \psi \cdot \boldsymbol{\xi} = \psi_{\boldsymbol{\xi}} = \lim_{t \to 0} \frac{\psi(\boldsymbol{x} + t\boldsymbol{\xi}) - \psi(\boldsymbol{x})}{t}$$

for $\psi$ considered as a function of $\boldsymbol{x}$.

**Theorem 1.** *Let $F : \mathbb{R}^n_+ \to \mathbb{R}^+$ be a $C^2$-function. Assume $F$ is strictly decreasing along open half lines in $\mathbb{R}^n_+$ that start from the origin, and maps such half lines surjectively to $\mathbb{R}^+$. Then the equation*

$$F\left(\frac{\boldsymbol{x}}{\psi}\right) = 1 \tag{23}$$

*defines* $\psi(\boldsymbol{x})$ *as an implicit 1-homogeneous positive function of* $\boldsymbol{x} \in \mathbb{R}^n_+$. *Setting* $\boldsymbol{x} = \psi(\boldsymbol{x})\boldsymbol{z}$, *the Hessians of* $F$ *and of* $\psi$ *are related by*

$$H_\psi(\boldsymbol{x})(\boldsymbol{\xi}, \boldsymbol{\eta}) = \frac{H_F(\boldsymbol{z})(\boldsymbol{\xi}, \boldsymbol{\eta})}{\psi(\boldsymbol{x})\langle DF(\boldsymbol{z}), \boldsymbol{z}\rangle}. \tag{24}$$

*Proof.* First we check that $\psi$ is 1-homogeneous in $\boldsymbol{x}$. If $\psi(\boldsymbol{x})$ is a solution of eq. (23) for a given $\boldsymbol{x}$, then $t\psi(\boldsymbol{x})$ is a solution for $t\boldsymbol{x}$, $t \in \mathbb{R}^+$, i.e., $\psi(t\boldsymbol{x}) = t\psi(\boldsymbol{x})$.

The condition for the implicit function theorem to apply to the function $\psi$ follows from formally differentiating (23) with respect to every $x_k$. Writing $F_i = F_i(\boldsymbol{z})$ for the partial derivative of $F$ with respect to $z_i$, and $\psi_k = \psi_k(x)$ for the partial derivative of $\psi$ with respect to $x_k$, we find

$$F_k(\boldsymbol{z})\,\frac{1}{\psi(\boldsymbol{x})} - F_i(\boldsymbol{z})\frac{x_i}{\psi(\boldsymbol{x})^2}\psi_k(\boldsymbol{x}) = 0,$$

in which we used the summation convention for $i$.

Multiplying by $\psi$ this reads

$$F_k = (z_iF_i)\psi_k, \tag{25}$$

and we see that the implicit function theorem condition is that $z_iF_i \neq 0$ in the point of consideration. It holds because $F$ is assumed to be strictly decreasing along lines through the origin, so

$$\langle DF(\boldsymbol{z}), \boldsymbol{z}\rangle = F_iz_i < 0.$$

We can then differentiate again, with respect to some $x_l$, to conclude that

$$F_{kl}\frac{1}{\psi} - F_{kj}\frac{x_j}{\psi^2}\psi_l = \left((z_iF_i)_l\frac{1}{\psi} - (z_iF_i)_j\frac{x_j}{\psi^2}\psi_l\right)\psi_k + (z_iF_i)\psi_{kl},$$

with summation convention for $i$ and $j$.

Multiplying by $\psi$ again we have

$$F_{kl} - F_{kj}\frac{x_j}{\psi}\psi_l = \left((z_iF_i)_l - (z_iF_i)_j\frac{x_j}{\psi}\psi_l\right)\psi_k, + \psi(z_iF_i)\psi_{kl}$$

or

$$\begin{aligned} F_{kl} - \underbrace{F_{kj}z_j\psi_l}_{z_jF_{jk}\psi_l} &= \left((z_iF_i)_l - (z_iF_i)_jz_j\psi_l\right)\psi_k + \psi(z_iF_i)\psi_{kl} \\ &= \psi_kF_l + \psi_kz_iF_{il} - \underbrace{\psi_kF_jz_j\psi_l}_{\psi_kF_l} - \psi_kz_iF_{ij}z_j\psi_l + (x_iF_i)\psi_{kl}, \end{aligned}$$

so

$$(x_iF_i)\psi_{kl} = F_{kl} + z_iz_jF_{ij}\psi_k\psi_l - z_jF_{jk}\psi_l - z_iF_{il}\psi_k.$$

Multiplying by $(z_iF_i)^2 = (z_jF_j)^2 = z_iz_jF_iF_j$ and using eq. (25) this yields

$$\begin{aligned} (z_iF_i)^3\psi\psi_{kl} &= z_iz_jF_iF_jF_{kl} + z_iz_jF_{ij}F_kF_l - z_iF_iz_jF_{jk}F_l - \underbrace{z_jF_jz_iF_{il}F_k}_{z_iF_iz_jF_{jl}F_k} \\ &= z_iz_j\left(F_iF_jF_{kl} + F_{ij}F_kF_l - F_iF_{jk}F_l - F_iF_{jl}F_k\right). \end{aligned}$$

Thus the bilinear form

$$\psi_{kl}\xi_k\eta_l,$$

with summation convention for $k$ and $l$, is a multiple of the bilinear form

$$\begin{aligned} &z_iz_j\left(F_iF_jF_{kl} + F_{ij}F_kF_l - F_iF_{jk}F_l - F_iF_{jl}F_k\right)\xi_k\eta_l \\ &\quad = (z_iF_i)(z_jF_j)(\xi_k\eta_lF_{kl}) + (z_iz_jF_{ij})(\xi_kF_k)(\eta_lF_l) \\ &\qquad - (z_iF_i)(z_j\xi_kF_{jk})(\eta_lF_l) - (z_iF_i)(z_j\eta_lF_{jl})(\xi_kF_k) \\ &\quad = \langle DF(\boldsymbol{z}), \boldsymbol{z}\rangle^2H_F(\boldsymbol{z})(\boldsymbol{\xi}, \boldsymbol{\eta}) + \langle DF(\boldsymbol{z}), \boldsymbol{\xi}\rangle\langle DF(\boldsymbol{z}), \boldsymbol{\eta}\rangle H_F(\boldsymbol{z})(\boldsymbol{z}, \boldsymbol{z}) \\ &\qquad - \langle DF(\boldsymbol{z}), \boldsymbol{z}\rangle\langle DF(\boldsymbol{z}), \boldsymbol{\eta}\rangle H_F(\boldsymbol{z})(\boldsymbol{z}, \boldsymbol{\xi}) \\ &\qquad - \langle DF(\boldsymbol{z}), \boldsymbol{z}\rangle\langle DF(\boldsymbol{z}), \boldsymbol{\xi}\rangle H_F(\boldsymbol{z})(\boldsymbol{z}, \boldsymbol{\eta}). \end{aligned} \tag{26}$$

Since $(z_iF_i)\psi = x_iF_i = \langle DF(\boldsymbol{z}), \boldsymbol{x}\rangle$, dividing eq. (26) by $(z_iF_i)^2 = \langle DF(\boldsymbol{z}), \boldsymbol{z}\rangle^2$ we get

$$
\begin{aligned}
&\langle DF(\boldsymbol{z}), \boldsymbol{x}\rangle \, H_\psi(\boldsymbol{x})(\boldsymbol{\xi}, \boldsymbol{\eta}) \\
&\qquad = H_F(\boldsymbol{z})(\boldsymbol{\xi}, \boldsymbol{\eta}) + \frac{\langle DF(\boldsymbol{z}), \boldsymbol{\xi}\rangle}{\langle DF(\boldsymbol{z}), \boldsymbol{z}\rangle} \frac{\langle DF(\boldsymbol{z}), \boldsymbol{\eta}\rangle}{\langle DF(\boldsymbol{z}), \boldsymbol{z}\rangle} H_F(\boldsymbol{z})(\boldsymbol{z}, \boldsymbol{z}) \\
&\qquad\quad - \frac{\langle DF(\boldsymbol{z}), \boldsymbol{\eta}\rangle}{\langle DF(\boldsymbol{z}), \boldsymbol{z}\rangle} H_F(\boldsymbol{z})(\boldsymbol{z}, \boldsymbol{\xi}) - \frac{\langle DF(\boldsymbol{z}), \boldsymbol{\xi}\rangle}{\langle DF(\boldsymbol{z}), \boldsymbol{z}\rangle} H_F(\boldsymbol{z})(\boldsymbol{z}, \boldsymbol{\eta}).
\end{aligned} \tag{27}
$$

Writing, using eq. (25),

$$
a = \frac{\langle DF(\boldsymbol{z}), \boldsymbol{\xi}\rangle}{\langle DF(\boldsymbol{z}), \boldsymbol{z}\rangle} = \langle D\psi(\boldsymbol{x}), \boldsymbol{\xi}\rangle, \quad b = \frac{\langle DF(\boldsymbol{z}), \boldsymbol{\eta}\rangle}{\langle DF(\boldsymbol{z}), \boldsymbol{z}\rangle} = \langle D\psi(\boldsymbol{x}), \boldsymbol{\eta}\rangle \tag{28}
$$

for the quotients of the components of $\boldsymbol{\xi}$ and $\boldsymbol{z}$ and $\boldsymbol{\eta}$ and $\boldsymbol{z}$ along $\nabla F(\boldsymbol{z})$, it follows that eq. (27) can be written as

$$
\langle DF(\boldsymbol{z}), \boldsymbol{x}\rangle \, H_\psi(\boldsymbol{x})(\boldsymbol{\xi}, \boldsymbol{\eta}) = H_F(\boldsymbol{z})(\boldsymbol{\xi} - a\boldsymbol{z}, \boldsymbol{\eta} - b\boldsymbol{z}). \tag{29}
$$

We also see that

$$
\begin{aligned}
\langle D\psi(\boldsymbol{x}), \boldsymbol{\xi}\rangle &= 0 = \langle D\psi(\boldsymbol{x}), \boldsymbol{\eta}\rangle \\
&\Longleftrightarrow \\
\langle DF(\boldsymbol{z}), \boldsymbol{\xi}\rangle &= 0 = \langle DF(\boldsymbol{z}), \boldsymbol{\eta}\rangle \\
&\Longrightarrow \\
H_\psi(\boldsymbol{x})(\boldsymbol{\xi}, \boldsymbol{\eta}) &= \frac{H_F(\boldsymbol{z})(\boldsymbol{\xi}, \boldsymbol{\eta})}{\langle DF(\boldsymbol{z}), \boldsymbol{x}\rangle} \\
&= \frac{H_F(\boldsymbol{z})(\boldsymbol{\xi}, \boldsymbol{\eta})}{\psi(\boldsymbol{x})\langle DF(\boldsymbol{z}), \boldsymbol{z}\rangle}.
\end{aligned} \tag{30}
$$

□

The preceding theorem concerns the Hessians acting as bilinear forms on vectors $\boldsymbol{\xi}$ and $\boldsymbol{\eta}$. For the quadratic form, when $\boldsymbol{\xi} = \boldsymbol{\eta}$, we can summarise eq. (27) as follows,

$$
\langle DF(\boldsymbol{z}), \boldsymbol{x}\rangle \underbrace{H_\psi(\boldsymbol{x})(\boldsymbol{\xi}, \boldsymbol{\xi})}_{\psi_{\boldsymbol{\xi}\boldsymbol{\xi}}} = H_F(\boldsymbol{z})(\boldsymbol{\xi} - \underbrace{\langle D\psi(\boldsymbol{x}), \boldsymbol{\xi}\rangle}_{\psi_{\boldsymbol{\xi}} = \nabla\psi\cdot\boldsymbol{\xi}} \boldsymbol{z}, \boldsymbol{\xi} - \langle D\psi(\boldsymbol{x}), \boldsymbol{\xi}\rangle \boldsymbol{z}), \tag{31}
$$

in which the $\boldsymbol{\xi}$-subscripts denote directional derivatives. In the same fashion (25) leads to

$$
F_{\boldsymbol{\xi}} = F_k\xi_k = \nabla F \cdot \boldsymbol{\xi} = (z_iF_i)\psi_k\xi_k = F_{\boldsymbol{z}}\psi_{\boldsymbol{\xi}} = (\nabla F \cdot \boldsymbol{z})(\nabla\psi \cdot \boldsymbol{\xi}). \tag{32}
$$

In particular eq. (25) implies

$$
\psi_{\boldsymbol{z}} = 1.
$$

Using subscripts eq. (25) and eq. (27) are best remembered as

$$
F_{\boldsymbol{z}}\psi_{\boldsymbol{\xi}} = F_{\boldsymbol{\xi}}, \quad F_{\boldsymbol{z}}\psi_{\boldsymbol{\xi}\boldsymbol{\xi}} = H_F(\boldsymbol{\xi} - \psi_{\boldsymbol{\xi}}\boldsymbol{z}, \boldsymbol{\xi} - \psi_{\boldsymbol{\xi}}\boldsymbol{z}), \tag{33}
$$

with $F$ and its derivatives evaluated in $\boldsymbol{z} = \frac{\boldsymbol{x}}{\psi(\boldsymbol{x})}$, and $\psi$, $\psi_{\boldsymbol{\xi}}$, $\psi_{\boldsymbol{\xi}\boldsymbol{\xi}}$ in $\boldsymbol{x}$.

**Corollary 2.** *If $H_F(\boldsymbol{z})$ is strictly positive definite and $\langle DF(\boldsymbol{z}), \boldsymbol{z}\rangle < 0$, then $\psi$ is strictly concave on $x_1 + \cdots + x_n = 1$.*

*Proof.* Since $\psi$ is 1-homogeneous, $H_\psi(\boldsymbol{x})(\boldsymbol{\xi}, \boldsymbol{\xi}) = 0$ if $\boldsymbol{\xi}$ is a multiple of $\boldsymbol{x}$. (In fact, it also holds that $H_\psi(\boldsymbol{x})(\boldsymbol{\xi}, \boldsymbol{\eta}) = 0$ if $\boldsymbol{\xi}$ and/or $\boldsymbol{\eta}$ is a multiple of $\boldsymbol{x}$.) But if $\boldsymbol{\xi}$ is not a multiple of $\boldsymbol{z}$, $H_\psi(\boldsymbol{x})(\boldsymbol{\xi}, \boldsymbol{\xi})$ cannot be zero. In particular, by Theorem 1 it is strictly negative definite on the affine set $x_1 + \cdots + x_n = 1$. □

1.4. **Concavity of specific flux for general EFMs.** To connect Theorem 1 to our EFM context, we need to understand what $\langle DF(\boldsymbol{z}), \boldsymbol{z}\rangle < 0$ and $\frac{\partial F}{\partial z_k} < 0$ for each $k = 1\dots, n$ exactly means in the context of an enzymatic pathway.

Recall that $\boldsymbol{z} = \boldsymbol{e}/J$. Points along a line through the origin in $\boldsymbol{z}$-space all have the same enzyme vector $\boldsymbol{e}$, but differ in flux (or, equivalently, they all have the same flux and same *relative* enzyme allocation, but the total enzyme concentration differs; we prefer to argue from the first standpoint). The term $\langle DF(\boldsymbol{z}), \boldsymbol{z}\rangle$ thus measures the change in nutrient concentration $\underline{x_0} = F(\boldsymbol{z})$ as $\boldsymbol{z}$ moves along this line through the origin. Hence

$$\langle DF(\boldsymbol{z}), \boldsymbol{z}\rangle < 0 \quad \Longleftrightarrow \quad \frac{dJ}{d\underline{x_0}}(\boldsymbol{e}) > 0. \tag{34}$$

The latter inequality is a natural property of metabolic pathways.

The term $\frac{\partial F}{\partial z_k}$ also has a natural interpretation.

**Lemma 3.** *Let $\underline{x_0} = F(\boldsymbol{z})$ be the metabolite steady state for an EFM pathway with $n$ reactions with nutrient concentration $\underline{x_0}$, and $\boldsymbol{z} = \boldsymbol{e}/J$. Then*

$$\langle DF(\boldsymbol{z}), \boldsymbol{z}\rangle \frac{\partial J}{\partial e_k} = \frac{\partial F}{\partial z_k}, \quad k = 1, \dots, n. \tag{35}$$

*Proof.* Differentiation of $\underline{x_0} = F(\boldsymbol{e}/J(\boldsymbol{e}))$ with respect to $e_k$ gives

$$\begin{aligned} 0 &= \sum_{j=1}^{n} \frac{\partial F}{\partial z_j}\frac{\partial z_j}{\partial e_k} \\ &= \sum_{j=1}^{n} \frac{\partial F}{\partial z_j}\left(\frac{1}{J}\delta_{jk} - \frac{e_j}{J^2}\frac{\partial J}{\partial e_k}\right). \end{aligned}$$

Therefore

$$\frac{\partial J}{\partial e_k}\sum_{j=1}^{n} e_j \frac{\partial F}{\partial z_j} = J\frac{\partial F}{\partial z_k}.$$

Substituting $e_j = Jz_j$ and using that $\langle DF(\boldsymbol{z}), \boldsymbol{z}\rangle = \sum_j z_j \frac{\partial F}{\partial z_j}$, the lemma follows. □

The term $\frac{\partial F}{\partial z_k}$ is therefore directly linked to $\frac{\partial J}{\partial e_k}$: under the condition $\langle DF(\boldsymbol{z}), \boldsymbol{z}\rangle < 0$, we have

$$\frac{\partial F}{\partial z_k} < 0 \quad \Longleftrightarrow \quad \frac{\partial J}{\partial e_k} > 0.$$

The assumption $\frac{\partial F}{\partial z_k} < 0$ is thus equivalent to the property that an increase of one enzyme concentration induces an increase in flux through the pathway.

We are now in a position to show that specific flux is strictly concave for a large class of EFM pathways.

**Theorem 4.** *Consider an EFM pathway of length $N$ with reaction rates $v_i = e_i f_i(\boldsymbol{x}, \underline{x_0})$, $i = 1\dots, N$ with $\underline{x_0}$ a nutrient input concentration. Let $J(\boldsymbol{e})$ be the flux through the pathway for some enzyme allocation $\boldsymbol{e}$ with total concentration $e_T$. Assume that*

1. *$Df(\boldsymbol{x}; \underline{x_0})$ is globally invertible;*
2. *an increase of $\underline{x_0}$ results in an increase of $J$, at fixed enzyme concentrations $\boldsymbol{e}$;*
3. *an increase in any enzyme concentration $e_k$ results in an increase of $J$, at fixed nutrient concentration $\underline{x_0}$;*
4. *the objective function eq. (18) is strictly convex.*

*Then $J(\boldsymbol{e})/e_T$ is strictly concave.*

*Proof.* By Section 1.3.1, the first assumption ensures $F(\boldsymbol{z})$ exists. By the second assumption, $\langle DF(\boldsymbol{z}), \boldsymbol{z}\rangle < 0$, see eq. (34). Together with the third assumption, Lemma 3 yields that $\frac{\partial F}{\partial z_k} < 0$ for all $k$. Thus, by Section 1.3.3, convexity of the objective function eq. (18) implies the convexity of $F(\boldsymbol{z})$. Theorem 1 and Corollary 2 now show that this is equivalent to concavity of $J/e_T$. □

1.5. **An explicit nontrivial example: the linear chain with Michaelis-Menten kinetics.** In this section we illustrate the techniques on the archetypal EFM, the linear chain. To keep the exposition clear, we limit ourselves to

$$\underline{x_0} \overset{e_1}{\rightleftharpoons} x_1 \overset{e_2}{\rightleftharpoons} x_2 \overset{e_3}{\rightleftharpoons} x_3 \overset{e_4}{\rightleftharpoons} \underline{x_4},$$

with $(e_1, e_2, e_3, e_4)$ in the enzyme simplex

$$\mathcal{E} \equiv \{(e_1, e_2, e_3, e_4) \in \mathbb{R}^4_{\geq 0} \mid e_1 + e_2 + e_3 + e_4 = e_T\}.$$

As rate laws we use the notation

$$f_i(x_{i-1}, x_i) = \frac{x_{i-1} - x_i}{a_i x_{i-1} + b_i x_i + c_i} \tag{36}$$

which may be related to the standard form

$$f_i = k_{cat,i} \frac{x_{i-1} - \frac{x_i}{K_{eq,i}}}{\frac{x_{i-1}}{K_{i,i-1}} + \frac{x_i}{K_{i,i}} + 1}$$

by scaling $x_i$ by $K_{eq,i}$ and setting

$$a_i = \frac{K_{eq,i}}{k_{cat,i} K_{i,i-1}}, \quad b_i = \frac{K^2_{eq,i}}{k_{cat,i} K_{i,i}}, \quad c_i = \frac{K_{eq,i}}{k_{cat,i}}.$$

The steady state equations are

$$e_i \frac{x_{i-1} - x_i}{a_i x_{i-1} + b_i x_i + c_i} = J, \quad i = 1, \ldots, 4. \tag{37}$$

Recall $x_0 = \underline{x_0}, x_4 = \underline{x_4}$ are given and fixed, take $\underline{x_0} > \underline{x_4}$ to have $J > 0$, and set $z_i = e_i/J$. Isolate $x_{i-1}$ as

$$x_{i-1} = \frac{z_i + b_i}{z_i - a_i} x_i + \frac{c_i}{z_i - a_i} \equiv \Phi_i(z_i, x_i) \tag{38}$$

to obtain

$$x_3 = F_4(z_4; \underline{x_4}), \quad x_2 = F_3(z_3, z_4; \underline{x_4}), \quad x_1 = F_2(z_2, z_3, z_4; \underline{x_4}),$$

and finally

$$\underline{x_0} = F_1(z_1, z_2, z_3, z_4; \underline{x_4}) \tag{39}$$

as the equation that defines the steady state solutions via the level sets of the family of functions $F_1$ parameterised by $\underline{x_4}$. To make this explicit, note that we have the recursion

$$x_{i-1} = \Phi_i(z_i, x_i) = \frac{c_i}{z_i - a_i} + \left(1 + \frac{a_i + b_i}{z_i - a_i}\right) \Phi_{i+1}(z_{i+1}, x_{i+1}),$$

so that

$$F_4(z_4; \underline{x_4}) = \frac{c_4}{z_4 - a_4} + \left(1 + \frac{a_4 + b_4}{z_4 - a_4}\right) \underline{x_4},$$

$$\begin{aligned} F_3(z_3, z_4; \underline{x_4}) &= \frac{c_3}{z_3 - a_3} + \left(1 + \frac{a_3 + b_3}{z_3 - a_3}\right) F_4(z_4; \underline{x_4}) \\ &= \frac{c_3}{z_3 - a_3} + \left(1 + \frac{a_3 + b_3}{z_3 - a_3}\right) \left(\frac{c_4}{z_4 - a_4} + \left(1 + \frac{a_4 + b_4}{z_4 - a_4}\right) \underline{x_4}\right) \\ &= \frac{c_3}{z_3 - a_3} + \frac{c_4}{z_4 - a_4}\left(1 + \frac{a_3 + b_3}{z_3 - a_3}\right) + \left(1 + \frac{a_3 + b_3}{z_3 - a_3}\right)\left(1 + \frac{a_4 + b_4}{z_4 - a_4}\right) \underline{x_4} \\ &= \gamma_3 M_3 + \gamma_4 M_4 (1 + M_3) + (1 + M_3)(1 + M_4)\, \underline{x_4}, \end{aligned}$$

in which

$$M_i = \frac{a_i + b_i}{z_i - a_i}, \quad \gamma_i = \frac{c_i}{a_i + b_i}. \tag{40}$$

Proceeding, we arrive at

$$\gamma_2 M_2 + \gamma_3 M_3 (1 + M_2) + \gamma_4 M_4 (1 + M_3)(1 + M_2) + (1 + M_2)(1 + M_3)(1 + M_4)\, \underline{x_4}$$

for $F_2(z_2, z_3, z_4; \underline{x_4})$. Finally we obtain the beautiful formula

$$\begin{aligned}
\underline{x_0} = F_1(z_1, z_2, z_3, z_4; \underline{x_4}) &= P_1(M_1, M_2, M_3, M_4; \underline{x_4}) \\
&= (1 + M_4)(1 + M_3)(1 + M_2)(1 + M_1)\,\underline{x_4} \\
&\quad +\gamma_4 M_4 (1 + M_3)(1 + M_2)(1 + M_1) \\
&\quad +\gamma_3 M_3 (1 + M_2)(1 + M_1) \\
&\quad +\gamma_2 M_2 (1 + M_1) \\
&\quad +\gamma_1 M_1
\end{aligned} \tag{41}$$

with natural domain $z_i > a_i$. Near the boundary of this domain, $F(\boldsymbol{z})$ grows unboundedly.

**Lemma 5.** *Let $F(\boldsymbol{z})$ be defined as in eq. (41), with $M_i$ and $\gamma_i$ as in eq. (40). Then $F$ decreases along lines through the origin,*

$$\langle DF(\boldsymbol{z}), \boldsymbol{z} \rangle < 0.$$

*Proof.* It may be checked from the explicit form eq. (41), with $M_i$ as in eq. (40), that $F$ decreases in each $z_k$, so it decreases along lines through the origin in the positive orthant. □

For the linear chain, convexity of $F(\boldsymbol{z})$ may be shown directly, as the following lemma shows.

**Lemma 6.** *Let $F(z)$ be defined as in eq. (41), with $M_i$ and $\gamma_i$ as in eq. (40). Then $F$ is strictly convex on the domain $z_i > a_i$, $i = 1, \dots, n$.*

*Proof.* Expressions like

$$g(t) = \frac{1}{(\zeta_1 - a_1 + \omega_1 t)(\zeta_2 - a_2 + \omega_2 t)}$$

differentiate as

$$g'(t) = -\left(\frac{\omega_1}{\zeta_1 - a_1 + \omega_1 t} + \frac{\omega_2}{\zeta_2 - a_2 + \omega_2 t}\right) g(t), \quad g''(t) = A(t) g(t) > 0,$$

because

$$A(t) = \left(\frac{\omega_1}{\zeta_1 - a_1 + \omega_1 t} + \frac{\omega_2}{\zeta_2 - a_2 + \omega_2 t}\right)^2 + \frac{\omega_1^2}{(\zeta_1 - a_1 + \omega_1 t)^2} + \frac{\omega_2^2}{(\zeta_2 - a_2 + \omega_2 t)^2}.$$

A second argument for strict convexity is to note that for $z_i > a_i$, we may divide each term in which $z_i$ appears by $z_i$, and expand using the geometric series. We immediately see that we obtain a sum of positive terms involving only powers of $a_i^n / z_i^n$, and that $F(\boldsymbol{z})$ is thus strictly convex.

As a third alternative, of course, since $F$ decreases in each $z_k$ variable and the objective function eq. (18) is strictly convex [4], the strict convexity of $F$ also follows from Section 1.3.3. □

Applying Theorem 1 we conclude that the Hessian of the implicitly defined function $J(e_1, e_2, e_3, e_4; \underline{x_0}, \underline{x_4})$ from

$$F\left(\frac{e_1}{J}, \frac{e_2}{J}, \frac{e_3}{J}, \frac{e_4}{J}; \underline{x_4}\right) = \underline{x_0}$$

is strictly negative definite in the $\boldsymbol{e}$-variables under the restriction that

$$e_1 + e_2 + e_3 + e_4 = e_T$$

is constant. That is, $J$, is strictly concave on the simplex defined by the latter constraint.

We could enlarge the scope of this direct approach of finding $F(\boldsymbol{z})$ explicitly somewhat by including all cases in which $c_i$ is replaced by an affine function of $x_{i+1}, \dots, \underline{x_4}$, i.e., including allosteric inhibition by products further down the line.

1.6. **A naive proof of concavity.** A naive but direct method to prove concavity is to show that the Hessian of $J(\boldsymbol{e}; \underline{x_0})$ is negative definite, using implicit differentiation on the quasi steady state equations. We give an example of this for a short chain. The method does not scale, and becomes infeasible for $n > 5$. Let $\mathcal{E} = \{\boldsymbol{e} \in \mathbb{R}^n_{\geq 0}$ such that $|\boldsymbol{e}| = 1\}$, and $\mathcal{X}_{\underline{x_0}} = \{\boldsymbol{x} \mid f_j(\boldsymbol{x}; \underline{x_0}) \geq 0\}$.

**Theorem 7.** *Consider $n = 3$. Then the map $J : \mathcal{E} \to \mathbb{R}$, $\boldsymbol{e} \mapsto J(\boldsymbol{e}; \underline{x_0})$ is strictly concave.*

*Proof.* The details of the computations that follow may be found in the supplementary Mathematica notebook. We prove that the determinant of the Hessian is strictly positive on $\mathcal{E}$. This shows that the product of the eigenvalues always has the same sign. A numerical check at one particular set of kinetic parameters then shows both eigenvalues are negative, making them always negative for all kinetic parameters. We first consider the QSS equations

$$e_1 f_1(\underline{x_0}, \bar{x}_1(e_1, e_2)) = J(e_1, e_2), \tag{42}$$

$$e_2 f_3(\bar{x}_1(e_1, e_2), \bar{x}_2(e_1, e_2)) = J(e_1, e_2), \tag{43}$$

$$(1 - e_1 - e_2) f_3(\bar{x}_2(e_1, e_2)) = J(e_1, e_2), \tag{44}$$

in which the QSS $(\bar{x}_1, \bar{x}_2)$ and $J$ are parameterised by $e_1$ and $e_2$, setting $e_3 = 1 - e_1 - e_2$. This parameterisation is warranted since the QSS is a smooth bijection between $\mathcal{E}$ and $\mathcal{X}_{\underline{x_0}}$. By implicit differentiation with respect to $e_1$ and $e_2$, and using the steady state equations eq. (42)–eq. (44), we find

$$\frac{\partial \bar{x}_1}{\partial e_1} = \frac{(f_2 f_3 + f_1 (f_2 + f_3))(f_3 (f_1 + f_3) f_{2,2} + f_1 f_2 f_{3,1})}{f_2 f_3 (f_3 f_{1,2} f_{2,2} + (f_2 f_{1,2} + f_1 f_{2,1}) f_{3,1})}, \tag{45}$$

$$\frac{\partial \bar{x}_1}{\partial e_2} = -\frac{(f_2 f_3 + f_1 (f_2 + f_3))\left(f_2^2 f_{3,1} - f_3^2 f_{2,2}\right)}{f_2 f_3 (f_3 f_{1,2} f_{2,2} + (f_2 f_{1,2} + f_1 f_{2,1}) f_{3,1})}, \tag{46}$$

$$\frac{\partial \bar{x}_2}{\partial e_1} = \frac{(f_2 f_3 + f_1 (f_2 + f_3))(f_2 f_3 f_{1,2} + f_1 (f_1 + f_3) f_{2,1})}{f_1 f_2 (f_3 f_{1,2} f_{2,2} + (f_2 f_{1,2} + f_1 f_{2,1}) f_{3,1})}, \tag{47}$$

$$\frac{\partial \bar{x}_2}{\partial e_2} = \frac{(f_2 f_3 + f_1 (f_2 + f_3))(f_2 (f_2 + f_3) f_{1,2} + f_1 f_3 f_{2,1})}{f_1 f_2 (f_3 f_{1,2} f_{2,2} + (f_2 f_{1,2} + f_1 f_{2,1}) f_{3,1})}, \tag{48}$$

$$\frac{\partial J}{\partial e_1} = \frac{f_1^2 f_{2,1} f_{3,1} - f_3^2 f_{1,2} f_{2,2}}{f_3 f_{1,2} f_{2,2} + (f_2 f_{1,2} + f_1 f_{2,1}) f_{3,1}}, \tag{49}$$

$$\frac{\partial J}{\partial e_1} = \frac{f_{1,2}\left(f_2^2 f_{3,1} - f_3^2 f_{2,2}\right)}{f_3 f_{1,2} f_{2,2} + (f_2 f_{1,2} + f_1 f_{2,1}) f_{3,1}}, \tag{50}$$

where we use the shorthand notation

$$f_i = f_i(\bar{x}_{i-1}, \bar{x}_i), \tag{51}$$

$$f_{i,j} = \frac{\partial f_i}{\partial x_j}(\bar{x}_{i-1}, \bar{x}_i) \tag{52}$$

for the partial derivatives, with $\bar{x}_0 = \underline{x_0}$ and $\bar{x}_3 = 0$; note the minus sign in the last expression, making all $f_{i,j} > 0$ on $\mathcal{X}_{\underline{x_0}}$.

Equations eq. (49) and eq. (50) provide us with explicit expressions of the dependence of $J$ on $\boldsymbol{e}$. We can differentiate them again with respect to the enzyme concentrations to find the second order partial derivatives,

$$\frac{\partial \bar{x}_i^2}{\partial e_j \partial e_k}, \qquad \frac{\partial J^2}{\partial e_j \partial e_k}, \qquad i, j, k = 1, 2.$$

Using the steady state equations again, and the first order derivatives eq. (45)–eq. (50) just computed, we can express $\frac{\partial J^2}{\partial e_j \partial e_k}$ completely into rational functions of $f_j$ and their first and second order partial derivatives. This leads to a concrete expression for the determinant of the Hessian of $J$,

$$\text{(53)}\quad \det\mathrm{H} = -\frac{(\mathrm{f}_2\mathrm{f}_3 + \mathrm{f}_1(\mathrm{f}_2+\mathrm{f}_3))^4}{\mathrm{f}_1\mathrm{f}_2\mathrm{f}_3(\mathrm{f}_3\mathrm{f}_{1,1}\mathrm{f}_{2,2} + (\mathrm{f}_1\mathrm{f}_{2,1} - \mathrm{f}_2\mathrm{f}_{1,1})\mathrm{f}_{3,2})^4}$$
$$f_{1,1}f_{3,2}\Big[2f_{2,2}\left(2f_1f_{1,1}f_{2,1}^2 + f_2\left(-2f_{2,1}f_{1,1}^2 - f_1f_{2,1,1}f_{1,1} + f_1f_{2,1}f_{1,1,1}\right)\right)f_{3,2}^2$$
$$+ f_3\Big\{2f_{2,1}\left(2f_{3,2}f_{2,2}^2 + f_2f_{3,2,2}f_{2,2} - f_2f_{3,2}f_{2,2,2}\right)f_{1,1}^2 +$$
$$f_1\left[2f_{3,2}f_{2,1,1}f_{2,2}^2\left(-2f_{3,2,2}f_{2,1}^2 - 4f_{3,2}f_{2,2,1}f_{2,1} + f_2f_{2,1,1}f_{3,2,2}\right)f_{2,2} +\right.$$
$$\left. f_{3,2}\left(2f_{2,2,2}f_{2,1}^2 + f_2\left(f_{2,2,1}^2 - f_{2,1,1}f_{2,2,2}\right)\right)\right]f_{1,1}$$
$$+ f_1f_{2,1}f_{1,1,1}\left(-2f_{3,2}f_{2,2}^2 - f_2f_{3,2,2}f_{2,2} + f_2f_{3,2}f_{2,2,2}\right)\Big\}\Big]$$

The notation for higher order derivatives here is

$$f_{i,j,k} = \frac{\partial^2 f_i}{\partial x_j \partial x_k}.$$

Note that this expression is in terms of $\underline{x_0}, x_1$ and $x_2$, but we have not computed the Hessian of $J$ with respect to $x_1$ and $x_2$.

Now we substitute the reaction kinetics

$$f_j(x_{j-1}, x_j) = \frac{x_{j-1} - k_jx_j}{a_jx_{j-1} + b_jx_j + c_j}, \quad j = 1, 2, 3,$$

where $x_0 = \underline{x_0}$, $x_1 = \bar{x}_1$ and $x_2 = \bar{x}_2$. Compute all the relevant derivatives, and set $s = \underline{x_0} - k_1x_1$, $t = x_1 - k_2x_2$ so that $s, t > 0$ for all $(\bar{x}_1, \bar{x}_2) \in \mathcal{X}_{\underline{x_0}}$. The determinant of the Hessian becomes a strictly positive rational function of (very many!) (positive) variables and (positive) parameters. □

#### 2. The adaptive control and its stability properties

In this section we introduce the adaptive control for enzyme synthesis rates. We reintroduce the ribosome concentration, so that EFMs have again $n$ enzymes and 1 ribosome. We show that the control induces a globally stable dynamical system, with all orbits converging to the unique steady state of maximal specific flux.

We assume that the EFM is in quasi steady state throughout, so that

$$e_if_i(\boldsymbol{x}; \underline{x_0}) = V_iJ, \quad i = 1, \dots, n, r.$$

As discussed in the previous section,

$$\frac{J}{e_T} = \frac{1}{\sum_{i=1}^{r} \frac{V_i}{f_i(\boldsymbol{x};\underline{x_0})}},$$

so that, with $z_i = e_i/J$,

$$|\boldsymbol{z}| \equiv z_1 + \cdots + z_n + z_r = \sum_{i=1}^{r} \frac{V_i}{f_i(\boldsymbol{x}; \underline{x_0})}.$$

Recall that $\lambda = \frac{e_rf_r(\boldsymbol{x};\underline{x_0})}{e_T} = \frac{J}{e_T}$. Hence (see also [1]),

$$\text{(54)}\quad \begin{aligned}\frac{1}{\lambda} &= \frac{e_1 + \cdots + e_n + e_r}{J} \\ &= \frac{\frac{JV_1}{f_1(\boldsymbol{x};\underline{x_0})} + \cdots + \frac{JV_n}{f_n(\boldsymbol{x};\underline{x_0})} + \frac{J}{f_r(\boldsymbol{x};\underline{x_0})}}{J} \\ &= \frac{V_1}{f_1(\boldsymbol{x};\underline{x_0})} + \cdots + \frac{V_n}{f_n(\boldsymbol{x};\underline{x_0})} + \frac{1}{f_r(\boldsymbol{x};\underline{x_0})},\end{aligned}$$

and the growth rate can be expressed as the harmonic mean of the rate laws, essentially,

$$\text{(55)}\quad \lambda = \frac{1}{\frac{V_1}{f_1(\boldsymbol{x};\underline{x_0})} + \cdots + \frac{V_n}{f_n(\boldsymbol{x};\underline{x_0})} + \frac{1}{f_r(\boldsymbol{x};\underline{x_0})}}.$$

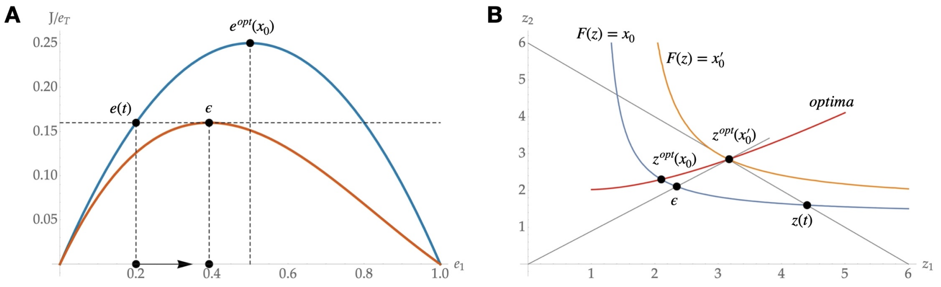


FIGURE 1. A schematic picture of the proof of global stability. A vector $\boldsymbol{e}$ is mapped to $F(\boldsymbol{z}) = \underline{x_0}$ by finding $J = J(\boldsymbol{e})$. Then to $\boldsymbol{z} = \boldsymbol{e}/J$ the point $\boldsymbol{w}$ is found, which is the unique optimum that lies on the same simplex as $\boldsymbol{z}$. Because level sets of $F$ are convex, and $F$ decreases along lines through the origin, $F(\boldsymbol{w}) < F(\boldsymbol{z})$. Now map $\boldsymbol{w}$ back to $F(\boldsymbol{z}) = \underline{x_0}$ by scaling, and call this point $\boldsymbol{e^{est}}$. Necessarily $\varepsilon_1 + \cdots + \varepsilon_n < z_1 + \cdots + z_n$, i.e., the specific flux at $\boldsymbol{\varepsilon}$ is higher than at $\boldsymbol{z}$. Since $\dot{\boldsymbol{e}} = \lambda(\boldsymbol{e^{est}} - \boldsymbol{e})$, $\boldsymbol{e}$ moves to a point with higher specific flux.

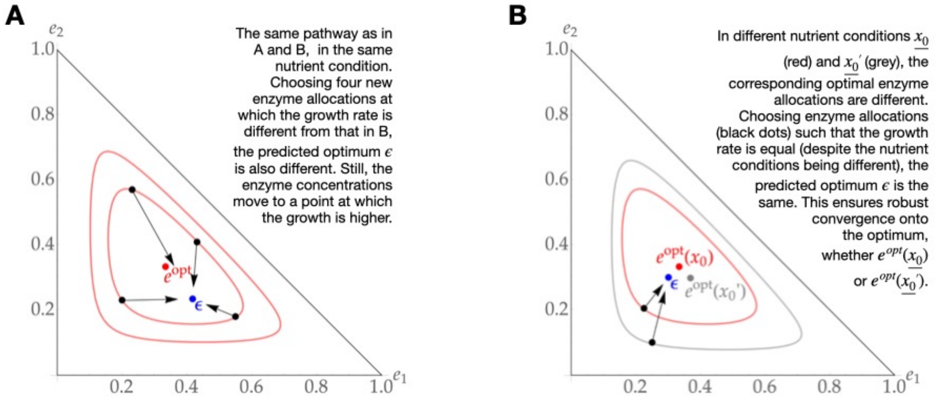


FIGURE 2. Additional situations indicating the robustness and global stability of qORAC control using growth rate as input. Compare with Figure 2 in the main text for details, and the captions in A and B above for further explanation.

All quasi steady states $\boldsymbol{x} = \boldsymbol{x}(\boldsymbol{e}; \underline{x_0})$ correspond to the surface $F(\boldsymbol{z}) = \underline{x_0}$ in $\boldsymbol{z}$-space; maximising $\lambda = J/e_T$ is thus equivalent to

$$\min \left\{ z_1 + \cdots + z_n + z_r \mid F(\boldsymbol{z}) = \underline{x_0} \right\}, \tag{56}$$

see Section 1.3.2. A simple Euler-Lagrange argument shows that any minimiser satisfies

$$D_1 F(\boldsymbol{z}) = \cdots = D_n F(\boldsymbol{z}) = D_r F(\boldsymbol{z}),$$

and $\underline{x_0}$ parametrises the curve of all optima.

The adaptive control now consists of predicting a pseudo-optimum at every time point, and allocating accordingly, see Figure 1 for an illustration. Its interpretation in terms of $J/e_T$ is Figure 3 in the main text.

**Lemma 8.** *Assume that $F(\boldsymbol{z})$ is strictly convex and decreasing along positive halflines through the origin. Consider for each $\underline{x_0}$ the minimiser of eq. (56). Then the set of optima is parametrised by $|\boldsymbol{z}| \equiv z_1 + \cdots + z_n + z_r$.*

*Proof.* Let $\underline{x_0}$ be given and $\boldsymbol{z}^{opt}(\underline{x_0})$ the corresponding unique optimum. Consider $\boldsymbol{z}(t) = (1-t)\boldsymbol{z}^{opt}$. Then $\boldsymbol{z}(t)$ lies on one fixed line through the origin, $\frac{d}{dt}F(\boldsymbol{z}(t))|_{t=0} > 0$ and $\frac{d}{dt}|\boldsymbol{z}(t)||_{t=0} < 0$. For $t > 0$, $\boldsymbol{z}(t)$ thus lies on a higher level set of $F$. On this level set, there exists a unique minimizer of eq. (56) again, $\boldsymbol{z}_t^{opt}$, say, and by convexity of $F$, $|\boldsymbol{z}_t^{opt}| \leq |\boldsymbol{z}(t)| < z^{opt}(\underline{x_0})$. The same argument holds for $t < 0$. □

**Corollary 9.** *Let $F$ be as in Lemma 8, and $\boldsymbol{z}$ such that $F(\boldsymbol{z}) = \underline{x_0}$ and $\boldsymbol{z}$ is not a minimiser of eq. (56). Then there exists $\xi_0 < \underline{x_0}$ such that $\boldsymbol{w}$ minimises $|\boldsymbol{w}|$ on $F(\boldsymbol{w}) = \xi_0$ such that $|\boldsymbol{z}| = |\boldsymbol{w}|$.*

*Proof.* Given any initial $\boldsymbol{z}$, $F(\boldsymbol{z})$ decreases as $\boldsymbol{z}$ moves outward along a straight line through the origin. In the mean time, $|\boldsymbol{z}|$ becomes unbounded. This is also true for minimisers. Hence there always exists an optimum with the required properties by decreasing $\xi_0$ sufficiently. □

Having found the pseudo-optimum $\boldsymbol{w}$, we allocate accordingly by setting

$$\chi_i = \frac{w_i}{|\boldsymbol{w}|},$$

so that $\sum_i \chi_j = 1$.

The adaptive control thus is given by

$$\begin{aligned} e_i f_i(\boldsymbol{x}; \underline{x_0}) &= V_i J \iff \underline{x_0} = F(\boldsymbol{z}), \\ \dot{e}_i &= e_r f_r(\boldsymbol{x}; \underline{x_0}) \chi_i - \lambda e_i, \\ \text{where } \chi_i &= \frac{w_i}{\sum_j w_j}, \\ D_1 F(\boldsymbol{w}) &= \cdots = D_n F(\boldsymbol{w}) = D_r F(\boldsymbol{w}) \\ \lambda &= \frac{J}{e_T} = \frac{1}{|\boldsymbol{w}|} = \frac{1}{|\boldsymbol{z}|}. \end{aligned} \tag{57}$$

We may also scale $\boldsymbol{w}$ back to the quasi-steady-state surface. Let $\boldsymbol{\varepsilon}$ be the multiple of $\boldsymbol{w}$ such that $F(\boldsymbol{e^{est}}) = \underline{x_0}$. In terms of $\boldsymbol{e^{est}}$, and using $\lambda = \frac{e_r f_r}{e_T}$ again, we may write the differential equation for $\dot{e}_i$ in eq. (57) as

$$\dot{e}_i = \lambda \left( \varepsilon_i - e_i \right). \tag{58}$$

We now have the crucial observation that the specific flux in the original pathway (so setting $x_0 = \underline{x_0}$) in $\boldsymbol{\varepsilon}$ is always higher than the current specific flux in $\boldsymbol{e}$.

**Corollary 10.** *Let $F$ be as in Lemma 8, and $\boldsymbol{z}$ such that $F(\boldsymbol{z}) = \underline{x_0}$ and $\boldsymbol{z}$ is not a minimiser of eq. (56). Let $\boldsymbol{w}$ be the predicted optimum such that $|\boldsymbol{w}| = |\boldsymbol{z}|$. Let $\boldsymbol{e^{est}}$ be the multiple of $\boldsymbol{w}$ such that $F(\boldsymbol{e^{est}}) = \underline{x_0}$. Then $|\boldsymbol{e^{est}}| < |\boldsymbol{z}|$.*

Let us summarise. Using $\boldsymbol{e}$ and the current specific flux $J$, a pseudo-optimal enzyme allocation with the same specific flux is predicted. This allocation corresponds to a pathway with lower nutrient concentration $\xi_0$ than the real pathway. If this enzyme allocation is plugged into the original pathway (by setting $x_0 = \underline{x_0}$ and letting the pathway come to a new quasi steady state), then the specific flux through this pathway is always higher than through the current pathway.

Since $J$ is strictly concave on this enzyme simplex, the $J$-level sets are strictly convex and nested. Hence $\boldsymbol{\varepsilon}$ lies strictly inside the flux level set defined by $\boldsymbol{e}$, and $\boldsymbol{e}$ moves towards it, see eq. (58). We conclude that $J$ increases along all orbits, and

$$\boldsymbol{e} \to \boldsymbol{e}^{opt}.$$

So in short,

**Theorem 11.** *Consider an EFM with dynamics specified by eq. (57). Then it is globally stable.*

A different way of writing the adaptive control uses the metabolite objective function

$$O(\boldsymbol{x}; \underline{x_0}) = \sum_j \frac{V_j}{f_j(\boldsymbol{x}; \underline{x_0})}$$

and states

$$
\begin{aligned}
e_i f_i(\boldsymbol{x}; \underline{x_0}) &= V_i J, \\
\dot{e}_i &= \varepsilon_i - \lambda e_i, \\
\varepsilon_i &= \frac{1/f_i(\boldsymbol{\xi})}{O(\boldsymbol{\xi}; \xi_0)} \\
O(\boldsymbol{\xi}; \xi_0) &= O(\boldsymbol{x}; \underline{x_0}) \\
\frac{\partial}{\partial \xi_k} O(\boldsymbol{\xi}; \xi_0) &= 0, \quad k = 1, \ldots, n, r.
\end{aligned} \tag{59}
$$

This form is very closely related to our previous adaptive control [4]. The only differences are that we have assumed that the EFM is in quasi-steady-state throughout, and the way in which a pseudo-optimum is predicted from the current metabolic state. Rather than using metabolite concentrations as sensors, we use the specific flux itself as sensor.

#### 3. Numerical simulations of qORAC

In this section we aim to show the reader how best to numerically explore qORAC-controlled elementary flux modes. Purely theoretically, a qORAC-controlled metabolic network is a described by a set of Differential-Algebraic Equations, or DAE. The differential equations describe only the enzyme and ribosome dynamics; the algebraic equations are of two kinds: the quasi steady state equations for metabolism, and the optimum equations with which the qORAC controller computes the predicted optimal ribosome allocation (also termed the optimal enzyme synthesis rates or optimal enzyme concentrations in different part of the paper).

The first and most important point that one must realise is that even though formally we should deal with DAEs, in practice one really shouldn't. The reason for this is that numerical integration of DAEs is hard. Rather than being able to start with arbitrary initial conditions, the solver needs to start with an initial condition in which many of the variables need to satisfy the algebraic equations. Finding such a starting condition is often as hard as the subsequent numerical integration. Moreover, once the starting condition has been checked by the solver to satisfy the algebraic equations sufficiently well, the solver needs to update these algebraic solutions in each time step, essentially by a numerical implementation of the Implicit Function Theorem. At the very least, this is very slow.

So rather than solving the DAE, it is much easier to

- treat the quasi steady state equations as the steady state of a dynamical system with metabolite concentrations as variables and enzyme/ribosome concentrations as parameters;
- Compute all optimal allocations before starting the simulations, and save them to disk, so they can be simply looked up from a table during the simulation.

We now provide a bit more detail on both steps.

3.1. **Remaining in quasi steady state.** At each time point, we assume that metabolism is in quasi steady state. We thus need $\boldsymbol{x}$ and $\boldsymbol{e}$ to satisfy

$$N\boldsymbol{v}(\boldsymbol{x}, \boldsymbol{e}; \underline{x_0}) = \mathbf{0} \tag{60}$$

for suitable model ingredients (the stoichiometric matrix $N$, rate laws that define the reaction rates $\boldsymbol{v}$). On the slow time scale, $\boldsymbol{e}$ changes by synthesis and dilution by growth; in the metabolic network, they are treated as parameters. So we should really view eq. (60) as

$$\dot{\boldsymbol{x}} = N\boldsymbol{v}(\boldsymbol{x}; \boldsymbol{e}, \underline{x_0}) \tag{61}$$

(Note that $\boldsymbol{e}$ appears now after the semicolon: it is now treated as fixed.) The initial condition for this ODE is the quasi steady state from the previous time step, and should be a good approximation to the one we are looking for in the current time step, as $\boldsymbol{e}$ has only changed a little in the mean time. Provided the metabolic network has unique steady states and is (at least locally) stable, convergence to the new quasi steady state should be very fast.

3.2. **Computing the optimal allocation.** At each time point, we need to prescribe an optimal ribosome allocation $\boldsymbol{\chi}$, or equivalently, optimal enzyme concentrations $\boldsymbol{\varepsilon}$. These should have the property, as qORAC dictates, that

(1) the corresponding growth rate is maximal for an environment with a nutrient concentration $\xi_0$ that is lower than the actual one $\underline{x_0}$;
(2) this maximal growth rate is equal to the current growth rate.

So for 1, $\boldsymbol{\varepsilon}$ should be a maximiser of

$$\max_{\boldsymbol{\xi};\boldsymbol{\varepsilon}}\{\lambda \mid JV_i = \varepsilon_j f_j(\boldsymbol{\xi};\xi_0),\ \lambda = J/\varepsilon_T, \varepsilon_j, f_j \geq 0,\ j = 1, \dots, n, r\}. \tag{62}$$

It is possible to characterise maximisers by a set of optimum equations. We have listed several versions in Section 2. For numerical computations, however, it is much easier to find the optimal allocations using a builtin routine, such as `fminsearch` in Matlab. Set up a map

$$\boldsymbol{e} \mapsto \lambda(\boldsymbol{e}; y_0)$$

that computes the growth rate of the metabolic network in quasi steady state at some fixed nutrient concentration $y_0$. Then let the optimisation routine vary $\boldsymbol{e}$ to maximise $\lambda$. By concavity of $\lambda(\boldsymbol{e}; y_0)$, this is also numerically a good problem. Once the optimum allocation has been found, write it to disk, including the optimal growth rate, optimal metabolite concentration and the nutrient concentration $y_0$ you prescribed.

Now vary $y_0$ from low to high and repeat the procedure. At the end you have a complete overview of all the optima (up to choosing the minimal and maximal nutrient concentration and the discretization of this interval of course). The optima are of course parameterised by the nutrient concentration, i.e., the map $x_0 \mapsto \boldsymbol{\varepsilon}(x_0)$ is well-defined. As shown in this paper, the optima can also be parameterised by $\lambda$ (a particular maximal growth rate only appears once in the table), and in many cases, also by internal metabolite concentrations (see [4]). The qORAC control in the numerical integration scripts now consists of nothing but a lookup from the table using linear interpolation on the chosen sensor (the growth rate in the case of this paper).



[1] Amsterdam Center for Dynamics and Computation, Department of Mathematics, VU University, 1081 HV Amsterdam, The Netherlands

[2] Systems Biology Lab, A-LIFE, AIMMS, VU University, 1081 HZ Amsterdam, The Netherlands
