## Supplementary material for "Sensing cellular growth rate facilitates its robust optimal adaptation to changing conditions": SI Matlab and Mathematica files: opt_egm.m

```
function alphaopt=opt_egm(x0);

alpha = [0.2;0.3;0.5];


MU = @(alpha) -qorac(10000,alpha,x0,0,1);

A = [1 1 1;
     -1 -1 -1;	
     -1 0 0 ;	 
     0 -1 0 ; 
     0 0 -1];
B = [1;-1;0;0;0];

opts = optimset('TolFun',1e-7);
[alphaopt] = fmincon(MU,alpha,A,B,[],[],[],[],[],opts)
```
