## Supplementary material for "Sensing cellular growth rate facilitates its robust optimal adaptation to changing conditions": SI Matlab and Mathematica files: qorac.m

```
function [mu,t,y,mus] = qorac(T,alpha,x0s,doplot,fixedalpha)

% function [mu,t,y,mus] = qorac(T,alpha,x0s,doplot,fixedalpha) 
%
% the qORAc script for qORAC control of the simple metabolic
% network shown in Figure 1 of 
% 
% Planqu\'e et al, Sensing cellular growth rate 
% facilitates robust optimal adaptation to changing conditions
% 
% See the Supplementary Information, section S3 for additional
% explanation of these simulations.
%
% Parameters:
% T: time span of one run (t in [0,T])
% alpha: ribosome allocation fractions 
%	alpha = 0 means alpha is not supplied. a default is chosen further down
%   alpha = 1 means we do qORAC control
%   alpha = a vector means we chose a fixed allocation. Used in finding optimal 
%			allocations by varying alpha
% x0s: one or more nutrient concentrations. When x0s is a vector, the simulation
% 	does a qORAC run from one nutrient concentration to the next, with the 
% 	network adapting to these changes
% doplot: flag for plotting (doplot=1, useful for individual runs) or not (doplot=0, useful 
% 			when finding optimal allocations, so this script is called many times in one run
%			run of scan_x0)
% fixedalpha is a flag with which we can change from finding optima 
% 	(in which we need to find the growth rate for fixed alphas all the time)
% 	to one in which we do qORAC control, changing the nutrient concentrations
% 	once every so often.
%   this flag is a separate one from the alpha parameter above, since alpha is sometimes 
% 	overwritten (it must become a vector at some point in the simulation).


% the optimal allocations (alphas, also termed epsilon in the paper, 
% since they differ up to a scalar constant)
% are computed using scan_x0.m
load('alphaopts1.mat')
load('muopts1.mat')
load('x0opts');

% the starting enzyme concentrations are not optimal. take something random away from optimal
% we use the alphas for that, but the alphas are the same as the enzyme concentrations,
% up to a constant. 
alpha0 = interp1(x0opts',alphaopt',x0s(1)) + 0.1*rand(1,3);
alpha0 = alpha0/(sum(alpha0));	% renormalise what you chose to sum = 1. 

% just some initial condition, using the alphas as enzyme concentrations
y0 = [0.2 0.4 alpha0(:)'];
% set up tolerances for the ODE solver, that are stricter than the built in defaults.
options = odeset('RelTol',1e-8,'AbsTol',1e-8);

% first we want to compute the quasi steady state without changing the enzymes
qssa=1;

% all parameters that are passed around in all functions below. 
% kcat1 and kcat2, kcat_ribo, current nutrient concentration x0, alpha, the optimal alphas and mu's, 
% and the qssa flag
pars = {[1,5], 10, x0s(1), alpha,alphaopt,muopts,qssa}; 

% the right hand side of the ODE for the QSSA system
QSSA = @(t,y) rhs(t,y,pars);
% pars 7 is the qssa flag. Set to 0 means we run 
% a network with enzyme dynamics (with or without qORAC control)
pars{7}=0;	
% this line seems the same as the QSSA line above, but pars has changed, so RHS is 
% really different from QSSA
RHS = @(t,y) rhs(t,y,pars);

% run the model to quasi steady state.
[tq,yq] = ode15s(QSSA,[0,1000],y0,options);
% take out the end point and make it the initial condition of the real simulation
y0 = yq(end,:);
% then run the real simulation
[t,y] = ode15s(RHS,[0,T],y0,options);


if fixedalpha~=1

  % for each nutrient concentration, we rerun the simulation above,
  % using the end of the previous run as the start of the next one
  for i=2:length(x0s)
    %update the nutrient concentration
    pars{3}=x0s(i);
    
    % set QSSA = 1, redefine the QSSA right hand side (note that x0 has changed inside pars)
    pars{7}=1;
    QSSA = @(t,y) rhs(t,y,pars);
    % same for the main model
    pars{7}=0;
    RHS = @(t,y) rhs(t,y,pars);

    % take the end of the last run as the start of the new one
    y0 = y(end,:);
    % run to QSS
    [tq,yq] = ode15s(QSSA,[0,1000],y0,options);
    y0 = yq(end,:);
    Tend = t(end);
    % run the next bit, and add it to the previous one
    [t2,y2] = ode15s(RHS,[Tend,Tend+T],y0,options);

    t = [t;t2];
    y = [y;y2];
  end
end
  
% the remaining part of the script is to compute statistics for all the 
% solutions, such growth rates   
mus = zeros(length(t),1);

for i=1:length(t)
  [~,~,mus(i)] = metab(y(i,:),pars);
end

% take away the semicolons if you wish to inspect these variables during output
alpha(:,:);
mu = mus(end);

if nargin < 3
  doplot=1;
end
  
% plot routines. turn off plotting by setting doplot=0  
if doplot
  figure(1)
  clf
  subplot(1,3,1)
  plot(t,y(:,1:2),'LineWidth',2)
  title('metab conc')

  subplot(1,3,2)
  plot(t,y(:,[3:5]),t,sum(y(:,[3:5]),2),'LineWidth',2)
  set(gca,'YLim',[0,1])
  legend('e1','e2','r1')
  title('enzymes and ribosomes')

  subplot(1,3,3)
  plot(t,mus,'LineWidth',2)
  legend('mu','Location','SouthEast')
  set(gca,'YLim',[0,1.2*max(mus(:))])
  title('growth rates')  
end

% save the results to disk
data = [t y mus];
save('qorac_example.txt','-ASCII','data');

%%%%%%%%%%%%%%%%%%%%%%%%%%%%%%%%%%%%%%%%%%%%%

function dy = rhs(t,y,pars);

% the main script in which the qORAC controlled network is defined


% get the current metabolite network state (fluxes, metabolite concentrations, growth rate)
[v,dx,mu] = metab(y,pars);

% pull out all the parameters
[kr,kc,x0,alpha,alphaopts1,muopts1,qssa] = pars{:};

% alpha is either 0 or 1. if alpha is zero, choose a particular vector
if alpha==0
  alpha = [0.2;0.3;0.5];

% if alpha is 1, do the qORAC control  
elseif alpha==1
  % as explained in the Supplementary Information, all optima are
  % precomputed, and we just pull out the one we want using linear interpolation on mu
  alpha = interp1(muopts1',alphaopts1',mu);
end  

% if we wish to run the script to compute a quasi steady state, the enzymes must be seen 
% as parameters, so their time derivatives must be zero.
if qssa == 1
  de1=0; de2=0; dr=0;
else
  e1 = y(3);  e2 = y(4);  r = y(5);

  % the enzyme equations are simply synthesis minus dilution. 
  de1 = v(3) * alpha(1) - mu * e1;
  de2 = v(3) * alpha(2) - mu * e2;
  dr = v(3) * alpha(3) - mu * r;
end

% collect all d/dt's into one vector
dy = [dx(:);de1;de2;dr];

%%%%%%%%%%%%%%%%%%%%%%%%%%%%%%%%%%%%%%%%%%%%%

function [v,dx,mu] = metab(y,pars);

% script to obtain the state of the metabolic network 
% we actually do not do the QSSA in each time step; the metabolite dynamics are
% much faster than the enzyme changes, anyway, so the difference between
% enforcing QSSA and not are minimal.

% pull out all the parameters
[kcs,kr,x0,alpha,alphaopts1,muopts1] = pars{:};

% the kcats of the enzymatic reaction
kc1=kcs(1);
kc2=kcs(2);

% the metabolite concentrations
x1 = y(1); x2 = y(2);
% the enzyme / ribosome concentrations
e1 = y(3); e2 = y(4); r = y(5);

% the reaction fluxes
v(1) = kc1 * e1 * (x0-x1) /(.3+2*x0+3*x1);
v(2) = kc2 * e2 * (x1-x2) / (1+x2+2*x1);
v(3) = kr * r * x2 / (.3+0.5*x2);

% total enzyme concentration
et = e1 + e2 + r;

% the growth rate is protein synthesis rate per unit protein
mu = v(3)  / et;

% the actual dynamical system for metabolism, with stoichiometry integrated.
dx(1) = v(1) - v(2) ;
dx(2) = v(2) - 10*v(3);
```
