## Supplementary material for "Sensing cellular growth rate facilitates its robust optimal adaptation to changing conditions": SI Matlab and Mathematica files: README

Instructions to reproduce simulations in 

	Sensing cellular growth rate facilitates its robust optimal adaptation to changing conditions

by 
	Robert Planqué, Josephus Hulshof and Frank J. Bruggeman

1. Put the contents of the zip file in a folder, open Matlab and go to the folder. 
2. Run in Matlab:
	>> scan_x0;
This will overwrite the files alphaopts1.mat, x0opts.mat, muopts1.mat
3. Run in Matlab:
	>> [mu,t,y,mus] = qorac(400,1,[.5,5,.2],1,0);
This reproduces the plots in the figure. The data is also exported to a textfile, which is 
read in with Mathematica, see the Matlab2Mathematica.nb notebook.

 
