## Supplementary material for "Sensing cellular growth rate facilitates its robust optimal adaptation to changing conditions": SI Matlab and Mathematica files: scan_x0.m

```
function [muopts,alphaopt]=scan_x0;

N=40;
x0opts=linspace(0.1,10,N)
for i=1:N
  x0 = x0opts(i);
  alphaopt(1:3,i) = opt_egm(x0);
  muopts(i) = qorac(10000,alphaopt(:,end),x0,0,1)
end

save('alphaopts1.mat','alphaopt');
save('muopts1.mat','muopts');
save('x0opts.mat','x0opts');

data = [x0opts(:) alphaopt', muopts(:)];
save('qorac_optima.txt','-ASCII','data');

figure(5)

subplot(1,2,1)
plot(muopts,alphaopt)
grid on
xlabel('growth rate')
ylabel('optimal allocations')
legend('e1','e2','r')

subplot(1,2,2)
plot(x0opts,alphaopt)
grid on
xlabel('nutrient concentration')
ylabel('optimal allocations')
legend('e1','e2','r')
```
